# Inhibition of the EBF1-ITGB8 Axis in Bone Marrow Niche Ameliorates Hallmarks of Myelofibrosis

**DOI:** 10.64898/2026.02.16.706171

**Authors:** Lyudmila Tsurkan, Mélodie Douté, Nicholas Morchel, Lahiri Konada, Rashid Mehmood, Te Ling, Amha Atakilit, Bridget Marcellino, Ronald Hoffman, Peter Vogel, Dean Sheppard, John D. Crispino, Marta Derecka

## Abstract

Fibrotic remodeling of the bone marrow (BM) niche is a characteristic feature of myelofibrosis (MF) that contributes to disease progression. In MF, mesenchymal stromal cells (MSCs) produce excessive amounts of inflammatory cytokines and extracellular matrix, leading to BM fibrosis, impaired blood production, extramedullary hematopoiesis, and progressive BM failure. While the genetic events that initiate MF in hematopoietic cells are well defined, our understanding of the mechanisms responsible for BM fibrosis remains incomplete. Here, we show that transcription factor EBF1 is a key regulator of the fibrotic gene program in mouse and human MSCs. *EBF1* is upregulated in pre-fibrotic MSCs, while mice with MSC-specific deletion of *Ebf1* exhibit reduced BM fibrosis, decreased expansion of myeloid cells and splenomegaly when transplanted with hematopoietic progenitors harboring the MF driver mutation MPL^W515L^. Moreover, we identify ITGB8 as an EBF1-regulated gene with therapeutic potential. MF mice treated with ITGB8-neutralizing antibodies or with MSC-specific *Itgb8* deletion show reduced disease burden, as indicated by decreased marrow fibrosis, significantly reduced frequencies of MPL mutant cells, and reduced inflammation in the BM. Our data indicate that targeting the EBF1-ITGB8 axis in the MF MSCs may have therapeutic benefits.

## Introduction

Hematopoietic stem cells (HSCs) and their malignant counterparts are maintained in a highly complex bone marrow (BM) microenvironment^1, 2^. Leptin receptor positive (LepR^+^) mesenchymal stromal cells (MSCs) are a crucial component of the BM niche, secreting critical growth factors necessary for maintenance of HSCs, myeloid and erythroid progenitors, but also essential for vascular and nerve regeneration^3–6^. However, in pathological hematopoiesis, aberrant blood cells disrupt the BM niche, including MSCs, which further impairs normal hematopoiesis and likely confers a competitive advantage to the mutant blood clones, facilitating disease evolution^7–11^.

Myelofibrosis (MF), the most aggressive form of myeloproliferative neoplasm (MPN), arises from somatic mutations in Janus kinase 2 (*JAK2*), thrombopoietin receptor (*MPL*), or calreticulin (*CALR*) acquired by HSC^12, 13^. These mutant blood cells induce maladaptive alterations in the BM niche via inflammatory cues and other factors such as transforming growth factor-beta (TGF-β) or platelet-derived growth factor (PDGF)^14^. Chronically inflamed MSCs in MF reduce the production of critical hematopoietic supportive factors like CXCL12 but produce an excessive amount of extracellular matrix (ECM), resulting in BM fibrosis and a defective microenvironment, which is thought to impair normal hematopoiesis and contribute to disease progression^8–10^. An unanswered question is how the same driver mutations generate three different clinical phenotypes of polycythemia vera (PV), essential thrombocythemia (ET), and MF. Interestingly, a recent study showed that spatially distinct BM niches influence the growth of *JAK2*^V617F^-mutated HSCs and clinical response to JAK inhibitor in PV vs ET, highlighting a key role for the HSC niche in explaining the variable pathogenesis and therapy response observed in different MPN subtypes^15^. MF patients suffer from initial expansion of myeloid cells, extramedullary hematopoiesis leading to hepatosplenomegaly, and progressive loss of hematopoiesis resulting in anemia and thrombocytopenia or transition to acute myeloid leukemia^13, 16^. JAK2 inhibitors, the current standard of care for MF patients, are effective in alleviating symptoms, decreasing inflammation and reducing spleen size^17^. However, they fail to eradicate mutant clones or attenuate BM fibrosis^16–18^. Moreover, many patients treated with JAK2 inhibitors become refractory or discontinue therapy due to side effects or loss of response to therapy, which results in poor outcomes^16, 18^. On the other hand, studies in mouse models have shown that targeting the defective fibrotic BM niche directly or by inhibiting aberrant communication between MSCs and mutant blood clones can halt MF progression^8, 9, 19–25^. Therefore, combining new treatments aimed at reducing BM fibrosis and sustaining BM microenvironment function with existing therapies directed against mutant hematopoietic clones might prove more effective for treating MF. Despite recent advances in mapping cellular composition and transcriptional perturbations in the BM niche during MF, the molecular programs underlying fibrotic remodeling, the major histopathological abnormality that occurs in MF, remain elusive. Thus, therapies preventing or reversing BM fibrosis remain an unmet need.

Here, we identified early B-cell factor 1 (EBF1) as a key transcriptional regulator of the fibrotic gene program in mouse and human MSCs during MF. EBF1 has been previously reported as one of the pivotal transcription factors necessary for MSC function, including differentiation towards mesenchymal lineages and HSC maintenance in the BM niche^26^. MSC-specific loss of EBF1 causes alterations in the cellular composition of the BM niche and impaired adhesion of HSCs, resulting in diminished myeloid output due to changes in the HSC’s chromatin landscape^26^. However, how EBF1 impacts MSC function in a disease context has not been explored. We show that EBF1 is upregulated in pre-fibrotic MSCs in response to hematopoietic cells with all three MF driver mutations, and that MSC-specific inactivation of *Ebf1* attenuates BM fibrosis in mouse models. Moreover, we identified integrin beta 8 (ITGB8) as an EBF1-regulated gene, which processes latent TGF-β to its active form^27–29^. In MF, TGF-β produced by inflamed MSCs and clonally expanded megakaryocytes and monocytes is a known driver of BM fibrosis^14, 23, 30^. Our results reveal that pharmacological inhibition of ITGB8 with neutralizing antibodies, as well as MSC-specific deletion of *Itgb8* reduces BM fibrosis and disease burden. Moreover, blocking ITGB8 results in reduced BM inflammation and improved BM niche function. Hence, we propose that targeting ITGB8 is an attractive approach to treat BM fibrosis and ameliorate MF.

## Results

### EBF1 is upregulated in mouse and human pre-fibrotic MSCs and fibroblasts

We have previously shown that transcription factor EBF1 is highly expressed in the mesenchymal compartment of the murine BM niche, where it plays a crucial role in HSC maintenance by regulating the expression of cytokines, adhesion molecules, integrins, and ECM components in MSCs^26^. We confirmed the expression of *Ebf1* in LepR^+^ and Cxcl12-expressing MSCs, as well as osteoblastic cells and arterial endothelial cells using an independent mouse single-cell RNA sequencing (scRNA-seq) dataset by Tikhonova et al. (Fig. 1a-c)^31^. Interestingly, genetic fate mapping and single-cell profiling studies identified LepR^+^ MSCs as the source of myofibroblasts producing the scar tissue in MF^8, 9, 22^. Considering that EBF1 is highly expressed in LepR^+^ MSCs and it controls the expression of many genes involved in BM fibrosis, including cytokines (*Il-1b*), alarmins (*S100a8* and *S100a9*), metalloproteinases, collagens, fibronectin and decorin (*Dcn*) in these cells, we hypothesized that EBF1 is a key regulator of the fibrotic remodeling of the BM niche in MF. First, we analyzed *Ebf1* expression in MSCs and fibroblasts using scRNA-seq datasets profiling the cellular landscape of the BM niche in MF mouse models^8, 19^. Analysis of the dataset from the Schneider group showed *Ebf1* expression in the majority of LepR^+^ MSCs isolated from mice transplanted with hematopoietic progenitors overexpressing thrombopoietin (ThPO), which induces BM fibrosis and MF phenotype (Extended Data Fig. 1a-b)^8^. Interestingly, *Ebf1* expression was enhanced in different subsets of BM niche cells from MF mice compared to the controls, (Extended Data Fig. 1c). Next, we leveraged scRNA-seq data from the Psaila group profiling the non-hematopoietic BM niche from mice transplanted with hematopoietic progenitors expressing the MPL^W515L^ mutation^19^. This analysis revealed abundant *Ebf1* expression in MSCs, fibroblasts, osteoblastic cells, pericytes, and endothelial cells (Fig. 1d-e). Moreover, *Ebf1* expression was significantly upregulated in fibrosis-driving fibroblasts in MF mice compared to the controls (Fig. 1e-f).

**Fig. 1.**
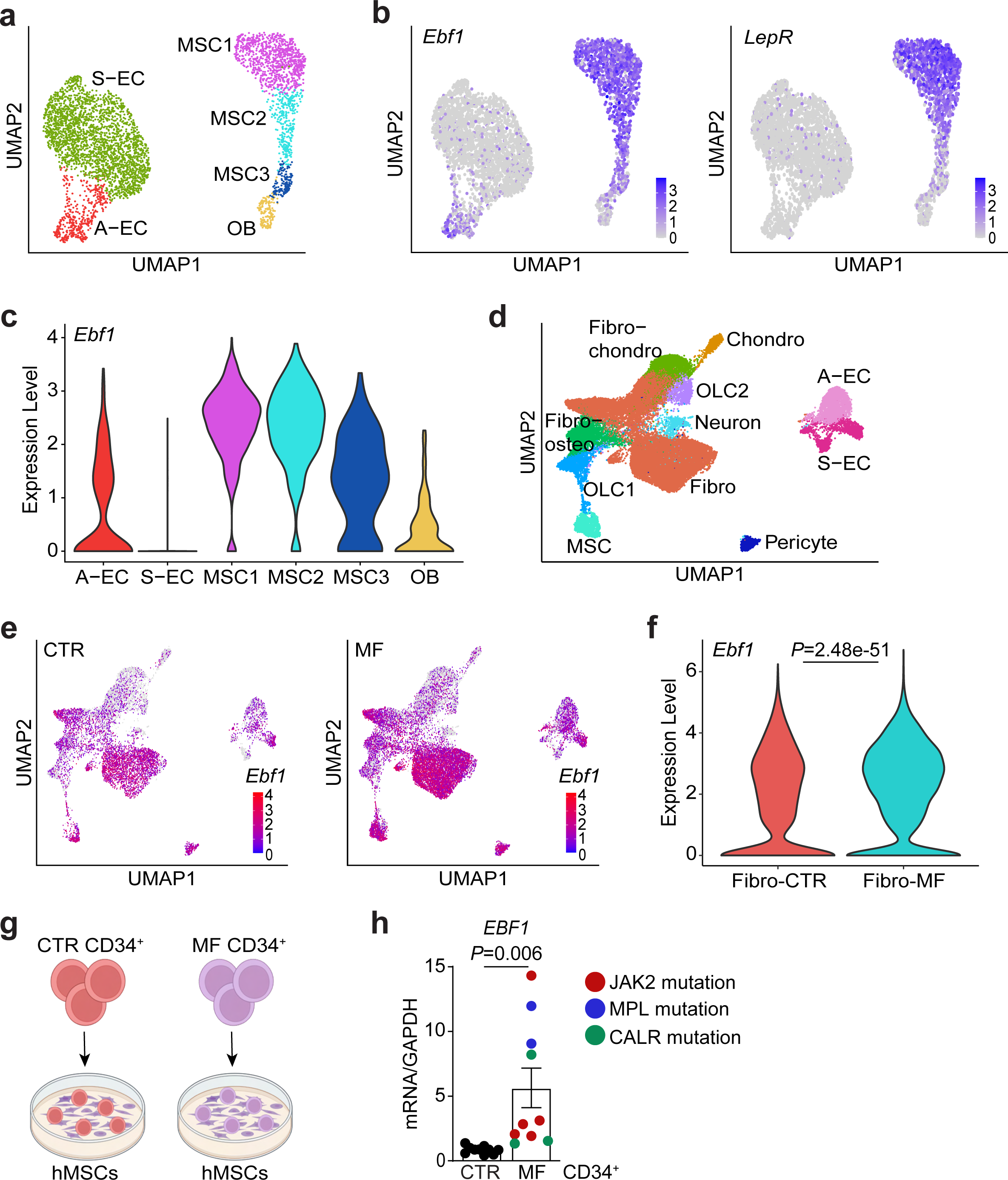
Increased *Ebf1* expression in MPN stroma. **a.** UMAP of non-hematopoietic BM niche cells isolated as reported by Tikhonova et al.; mesenchymal stromal cells (MSCs); osteoblastic cells (OB); sinusoidal endothelial cells (S-EC); arterial endothelial cells (A-EC). **b.** UMAP of normalized *Ebf1* and *LepR* expression in MSC clusters shown in (a). **c.** Violin plot showing normalized *Ebf1* expression in non-hematopoietic BM niche cells cluster identified in (a). **d.** UMAP of BM niche cells isolated from wild-type mice transplanted with hematopoietic progenitors expressing GFP (CTR) or MPL^W515L^ (MF) as described by Li, Colombo, Wang et al.; mesenchymal stromal cells (MSCs); osteolineage cells (OLC); fibroblasts (Fibro); neuronal cells (Neuron); chondrocytes (Chondro); sinusoidal endothelial cells (S-EC); arterial endothelial cells (A-EC). **e.** Normalized *Ebf1* expression in BM niche cells identified in (d) between control and MF mice. **f.** Violin plot showing increased *Ebf1* expression in fibroblast cluster shown in (d) and (e) from MF mice compared to the CTR. **g.** Experimental setup. **h.** *EBF1* expression in human BM MSCs co-cultured for 72h with CD34^+^ hematopoietic progenitors purified from the peripheral blood of healthy donors (Ctr CD34^+^) or MF patients (MF CD34^+^) harboring mutations in *JAK2* (5 patients; red dots), *MPL* (2 patients; blue dots) and *CALR* (3 patients; green dots). Data are the mean fold change ± SEM over the control; n=6 independent donors for MSCs; n=10 age and gender-matched donors of Ctr CD34^+^ cells and n=10 donors of MF CD34^+^ cells.

To determine whether the regulation of *EBF1* expression is conserved between mouse and human cells, we co-cultured human BM-derived MSCs (hMSCs) with CD34^+^ cells collected from the peripheral blood (PB) of MF patients or age-matched healthy donors (Fig. 1g). Similarly to mouse models of MF, *EBF1* expression was also upregulated in hMSCs co-cultured for 72 hours with hematopoietic progenitors harboring mutations in *JAK2* or *MPL* or *CALR* genes compared to the co-cultures with healthy hematopoietic progenitors (Fig. 1h). This is consistent with the presence of BM fibrosis in MF patients harboring any of the three driver mutations. Together, our analysis confirms increased *EBF1* expression in BM niche cells in response to hematopoietic cells carrying MF mutations. More importantly, this response is conserved in mouse and human and across different MF mouse models and MF driver mutations, suggesting that EBF1 orchestrates a broad transcriptional program driving fibrotic remodeling of the BM niche in MF.

### *Ebf1* deficiency in MSCs protects from BM fibrosis and MF progression

Based on these results, we reasoned that loss of *Ebf1* in MSCs might reduce BM fibrosis in MF. To investigate this, we used a well-established mouse model based on retroviral transduction of c-kit^+^ hematopoietic progenitors with the MPL^W515L^ (GFP^+^) variant identified in MPN patients^32^. It is also a clinically relevant model since the recipients develop hallmarks of human myeloproliferative disease, such as expansion of myeloid cells, increased megakaryocytes and thrombocytosis, extramedullary hematopoiesis with splenomegaly, decreased BM cellularity, and BM fibrosis^32^. We transplanted MPL^W515L^ -expressing cells into sub-lethally irradiated recipients with MSC-specific *Ebf1* deletion (*Prx1*^Cre^*Ebf1*^fl/fl^) or littermate controls (*Prx1*^Cre^*Ebf1*^+/+^ or *Ebf1*^fl/fl^) (Extended Fig. 2a). Flow cytometric analysis at the end point (4-5 weeks post-transplant) revealed decreased frequency of GFP^+^ MPL^W515L^ cells in the BM, but also spleen and peripheral blood of *Prx1*^Cre^*Ebf1*^fl/fl^ recipients compared to the controls (Extended Data Fig. 2b). Also, mice with *Ebf1*-deficient niche had diminished splenomegaly (Extended Data Fig. 2c). Most importantly, reticulin staining of BM sections showed significantly reduced fibrosis in the recipients with MSC-specific *Ebf1* deletion compared to the controls (Extended Data Fig. 2d).

Since *Prx1^Cre^* driven recombination occurs in a wide range of BM MSCs and osteoblastic cells, we extended our analysis to a more defined *LepR^Cre^* model to delete *Ebf1* in fibrosis-producing LepR^+^ MSCs (Fig. 2a). Analogously to *Prx1^Cre^*model, transplantation of MPL^W515L^ cells into *LepR*^Cre^*Ebf1*^fl/fl^ mice resulted in reduced frequency of MPL mutant cells in the BM, spleen and peripheral blood compared to the control mice (*LepR*^Cre^*Ebf1*^+/+^ or *Ebf1*^fl/fl^) (Fig. 2b). The reduced disease burden was mostly due to decreased expansion of mutant granulocytes (Mac1Gr1^hi^-GFP^+^) and monocytes (Mac1Gr1^lo^-GFP^+^) in the examined tissues (Extended Data Fig. 2e-h). Moreover, *LepR*^Cre^*Ebf1*^fl/fl^ recipients exhibited decreased spleen weights (Fig. 2c) and significantly reduced BM fibrosis based on reticulin staining (Fig. 2d and Extended Data Fig. 2i). These results confirm a key role for EBF1 in fibrotic remodeling of the BM niche and suggest that targeting the EBF1-controlled transcriptional program in the MSCs could ameliorate BM fibrosis and consequently MF progression.

**Fig. 2.**
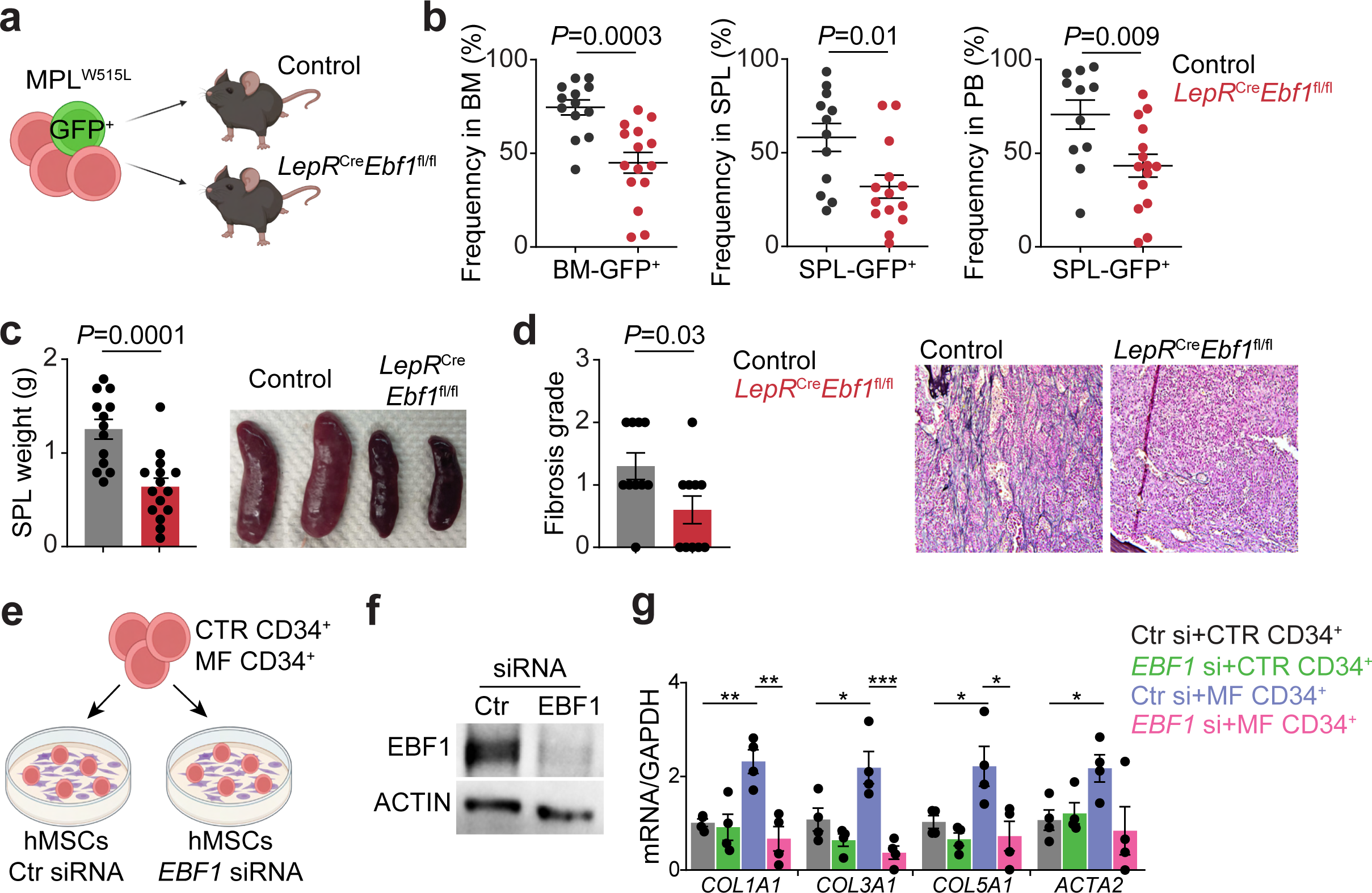
*Ebf1*-deficient BM niche reduces MF progression. **a.** Experimental setup. **b.** Frequency of MPL^W515L^ mutant (GFP^+^) cells in bone marrow (BM), spleen (SPL) and peripheral blood (PB) of the control and *LepR*^Cre^*Ebf1*^fl/fl^ recipient mice transplanted with MPL^W515L^ expressing cells. **c.** Spleen weights and representative image of spleens of the recipient mice. Each dot represents an individual recipient, data represent the mean ± SEM, *n*=11-13 controls and *n*=14-15 *LepR*^Cre^*Ebf1*^fl/fl^ mice in 3 independent experiments. **d.** Fibrosis grade scored based on reticulin staining and representative images of reticulin staining in BM sections (40X objective) of mice transplanted with MPL^W515L^ expressing cells. Each dot represents an individual recipient, data represent the mean ± SEM, *n*=10 controls and *n*=10 *LepR*^Cre^*Ebf1*^fl/fl^ mice. **e.** Experimental setup. **f.** EBF1 protein in human BM MSCs, 48 h after siRNA-mediated knockdown. **g.** Relative mRNA levels of fibrotic markers in control and *EBF1*-knockdown human MSCs co-cultured for 72h with CD34^+^ progenitors from the blood of healthy donors (CTR CD34^+^) or MF patients (MF CD34^+^) harboring JAK2^V617F^mutation. Co-cultures were initiated 48h after siRNA treatments. Data are the mean fold change ± SEM over the control; n=3 donors for MSCs and n=4 donors for healthy and MF CD34^+^ cells *p ≤0.05, **p ≤0.01, ***p ≤0.001.

### EBF1 regulates a fibrotic gene program in hMSCs

Given that *EBF1* upregulation in fibrosis-driving MSCs is conserved between mouse and human cells, we examined the impact of *EBF1* deletion on the activation of fibrotic markers in hMSCs co-cultured with CD34^+^ cells from MF patients (MF CD34^+^) or healthy donors (CTR CD34^+^) (Fig. 2e). We performed siRNA-mediated *EBF1* knockdown in the hMSCs prior to co-cultures and verified the knockdown using immunoblot analysis (Fig. 2f). hMSCs with non-targeting siRNA (Ctr si) co-cultured with CD34^+^ cells from MF patients showed increased expression of collagens and alpha-smooth muscle actin (encoded by *ACTA2* gene), the typical fibrotic markers, relative to co-cultures with healthy CD34^+^ cells (Fig. 2g). Collagen deposition in BM, in addition to reticulin, is part of the diagnostic criteria used to assess the degree of BM fibrosis in MF patients^33^. In contrast, the expression of fibrosis-related genes such as *COL1A1*, *COL3A1*, *COL5A1*, and *ACTA2* was blunted in *EBF1*-deficient hMSCs co-cultured with MF CD34^+^ cells compared to the controls (Fig. 2g). These data confirm that EBF1 activates a fibrotic gene program in mouse and human MSCs during MF.

### EBF1 occupies chromatin regions within genes contributing to fibrosis in MSCs

To gain further mechanistic insight into EBF1-dependend regulation of the fibrotic gene program, we performed Cleavage Under Targets & Release Using Nuclease (CUT&RUN) assay to investigate EBF1 chromatin binding. We isolated and expanded MSCs from wild-type C57BL/6J mice prior to nuclei isolation and subsequent immunoprecipitation with anti-EBF1 antibodies or control IgG (Fig. 3a). We detected substantial and highly specific EBF1 chromatin occupancy compared to IgG controls and confirmed concordance among the biological replicates with 1087 EBF1 peaks common between all replicates (qvalue cutoff = 1.00e-02) (Fig. 3b and Extended Data Fig. 3a). On a genome wide scale, EBF1 binds mostly intergenic and intronic regions, which are populated by regulatory elements like enhancers. (Fig. 3c and Extended Data Fig. 3b). Indeed, it has been shown that primary EBF1 binding regions in B cells are regulatory elements^34, 35^. Moreover, 5-7% of detected EBF1 peaks are localized to transcription start sites (TSS) within the promoter regions in MSCs (Fig. 3c and Extended Data Fig. 3b). Most importantly, EBF1 occupancy was enriched at the promoter regions of multiple fibrotic markers such as *Col3a1*, *Col5a1*, *Acta2*, *Dcn*, *Tgfb1* and within the *Col1a1* locus (Fig. 3d and Extended Data Fig. 3c). Motif analysis confirmed that EBF-specific binding sequences were the highest enriched motifs validating the specificity of our CUT&RUN data set (Fig. 3e). Interestingly, motif analysis revealed also an enrichment of AP-1 motifs within EBF1-bound regions in MSCs (Fig. 3e). Transcription factors from AP-1 family have been associated with tissue fibrosis. Elevated levels of c-JUN have been shown in biopsies of patients with various fibrotic diseases, including MF, and overexpression of c-JUN and JunB in mice leads to fibrosis in multiple organs^36^. Also, Fra-2 has been described as an important regulator of collagen production and osteoblast differentiation^37^. Overexpression of Fra-2 in mice results in osteosclerotic phenotype, a feature of advanced MF^37, 38^. It is reasonable to speculate that EBF1 and AP-1 factors cooperate to drive fibrotic phenotype in the BM. Our results indicate that EBF1 regulates the expression of the fibrotic genes via direct binding within their DNA regulatory regions.

**Fig. 3.**
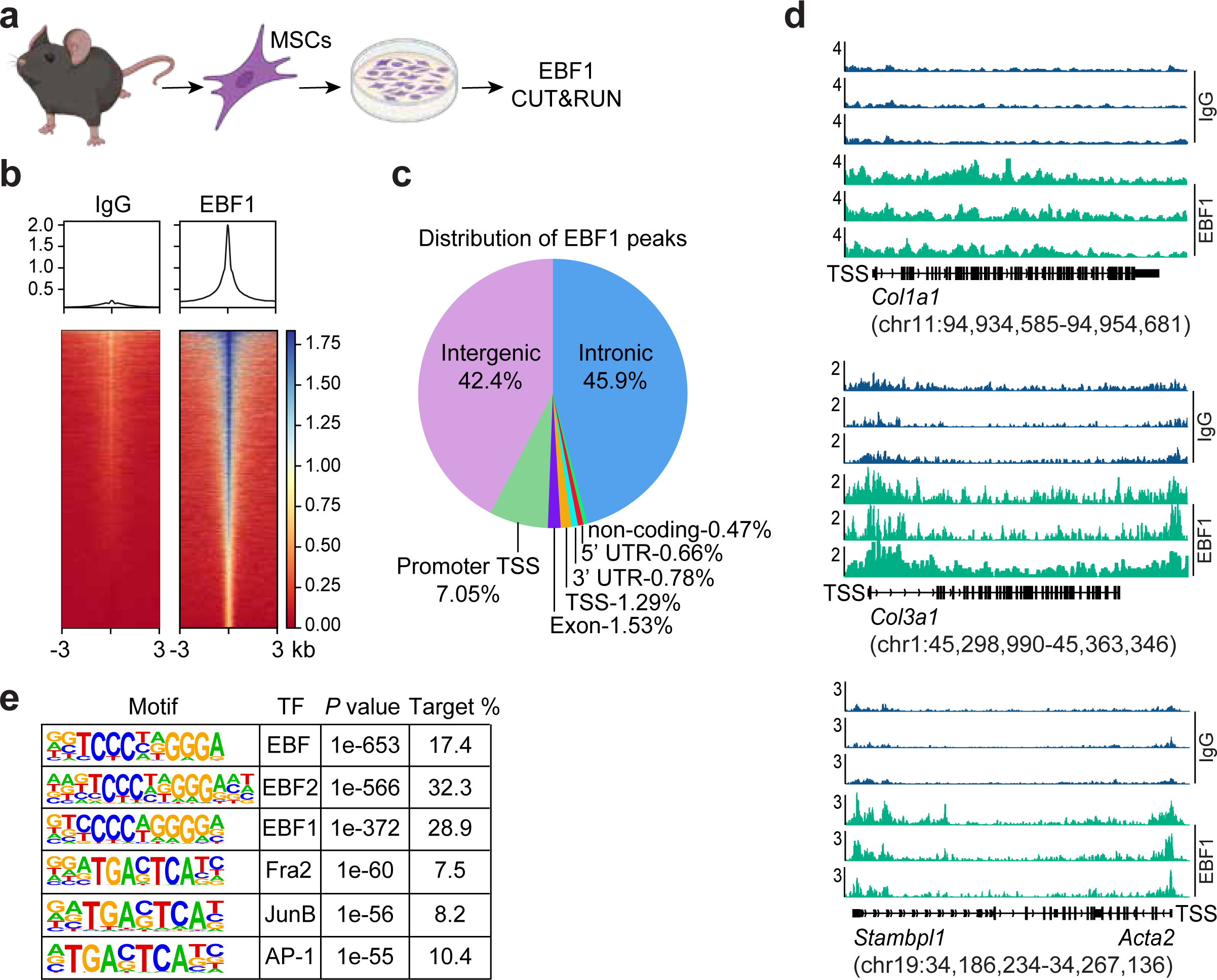
EBF1 binds within chromatin regions of fibrotic genes. **a.** Experimental design. **b.** Density plot showing chromatin binding of EBF1 in MSCs isolated from wild-type mice. EBF1 binding profile was normalized to control IgGs. Regions ±3 kb around EBF1 or IgG peak summit are represented as density maps with the average read coverage profile shown above the density maps. **c.** Percentage of EBF1 peaks in different genomic regions in mouse MSCs. **d.** Representative tracks showing EBF1 occupancy within *Col1a1*, *Col3a1* and *Acta2* loci in mouse MSCs (three independent biological replicates). The scale on the *y* axis represents RPKM values; transcriptional start site (TSS). **e.** DNA motif enrichment analysis within EBF1 CUT&RUN peaks.

### EBF1 controls the expression of pro-fibrotic ITGB8

The fact that EBF1 deficiency in MSCs renders them insensitive to fibrotic remodeling in MF suggests that reducing EBF1 activity in the BM niche could have therapeutic benefits. However, targeting EBF1 itself poses challenges due to its indispensable role in MSCs, which in turn are necessary for HSC maintenance, and its critical function in B cell development^26, 34^. Therefore, we turned to EBF1-regulated genes as potential therapeutic targets to ameliorate BM fibrosis. We have previously reported that EBF1 regulates expression of ECM genes in MSCs and transcriptome analysis of MSCs isolated from control and *Prx1*^Cre^*Ebf1*^fl/fl^ mice in that study identified integrin β8 (ITGB8) as one of the significantly downregulated genes in *Ebf1*-deficient MSCs^26^. Interestingly, ITGB8 has been implicated in solid organ fibrosis since its major function is processing latent (inactive) TGF-β to an active form, which promotes tissue fibrosis in BM, lungs, heart, kidney, and liver downstream of the TGF-β receptor^13, 29, 39, 40^. Remarkably, the bioavailability and pro-fibrotic function of TGF-β in the tissues is regulated mostly via its activation rather than its synthesis or secretion^41^. In MF, latent TGF-β is produced by multiple cell types, including expanded megakaryocytes and monocytes, as well as mast cells and MSCs^19, 23^. However, the cellular entities and mechanisms responsible for TGF-β activation and subsequent fibrosis are not completely clear. We propose that EBF1-regulated ITGB8 expression in MSCs contributes to fibrotic remodeling in the context of MF. Using CUT&RUN analysis, we confirmed EBF1 occupancy within the ITGB8 promoter and intronic regions, indicating that EBF1 is a direct transcriptional regulator of *Itgb8* expression (Fig. 4a). To further validate this direct regulation, we performed siRNA-mediated knockdown of *EBF1* in mouse and human MSCs, which corroborated reduced ITGB8 expression in both species within 48 hours upon EBF1 deletion (Fig. 4b-c).

**Fig. 4.**
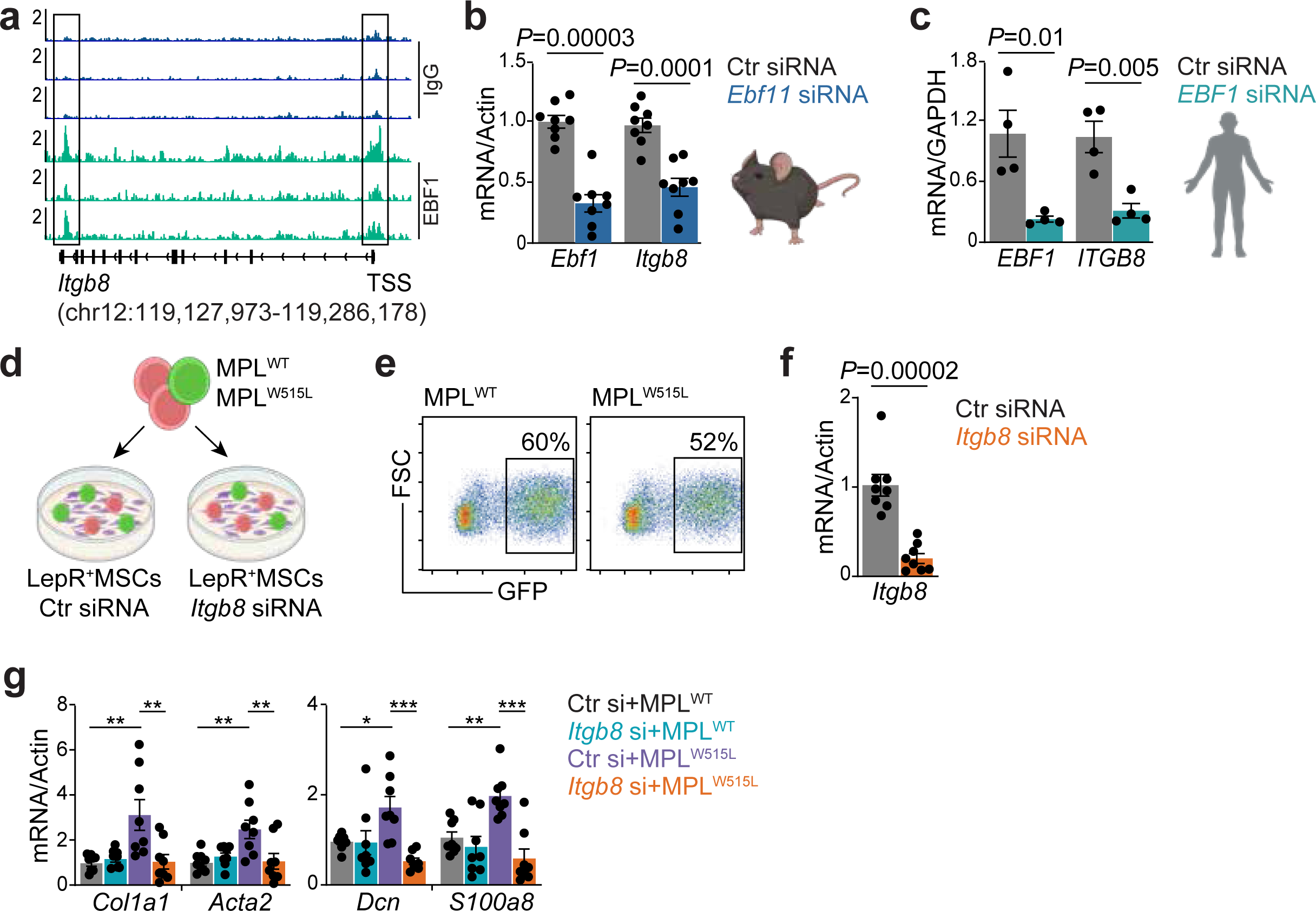
EBF1 regulates *ITGB8* expression in mouse and human MSCs. **a.** Genome tracks showing Cut&Run analysis assessing chromatin occupancy of EBF1 within ITGB8 locus in mouse MSCs. **b.** Relative *Ebf1* and *Itgb8* expression in mouse LepR^+^MSCs 72h after siRNA mediated *Ebf1* knockdown. Data are the mean fold change ± SEM over the control siRNA, each dot represents an individual mouse, n=8 wild-type mice. **c.** Relative *EBF1* and *ITGB8* expression in human MSCs 72h after siRNA mediated *EBF1* knockdown. Data are the mean fold change ± SEM over the control siRNA, each dot represents an individual healthy donor, n=4 donors. **d.** Experimental schematics. **e.** Representative flow cytometry plots showing efficiency of transduction with MPL^WT^ and MPL^W515L^ based on GFP frequency within c-kit^+^ cells used for co-cultures. **f.** Relative *Itgb8* expression in mouse LepR^+^MSCs 96h after siRNA mediated *Itgb8* knockdown (end point of co-cultures). Data are the mean fold change ± SEM over the control siRNA, each dot represents LepR^+^ MSCs from an individual mouse. **g.** Relative mRNA levels of fibrotic markers in control (Ctr si) and *Itgb8*-knockdown (*Itgb8* si) LepR^+^ MSCs co-cultured for 72h with either MPL^WT^ or MPL^W515L^ expressing cells. Co-cultures were initiated 24h after *Itgb8* knockdown in LepR^+^ MSCs. LepR^+^ MSCs were isolated from n=8 individual wild-type mice represented by each dot. Data are the mean fold change ± SEM over the control siRNA.

To evaluate whether ITGB8 contributes to the activation of fibrotic gene program in MSCs, we utilized co-cultures of LepR^+^ MSCs with *Itgb8* knockdown and c-kit^+^ hematopoietic progenitors expressing MPL^WT^ or MPL^W515L^ (Fig. 4d-f). Indeed, *Itgb8*-deficient LepR^+^ MSCs showed significantly reduced expression of fibrotic markers such as *Col1a1*, *Acta2*, and *Dcn* in response to MPL mutant cells relative to control LepR^+^ MSCs (Fig. 4g). Moreover, MSC-specific *Itgb8* knockdown resulted in lower expression of proinflammatory mediator *S100a8*, which has been recently described as a marker of disease progression toward the fibrotic stage in MPN mouse models and patients (Fig. 4g)^8^. Together, these results indicate that EBF1 and its target ITGB8 regulate the expression of genes contributing to BM fibrosis.

### MSC-specific loss of *Itgb8* attenuates MF development

To explore whether MSC-specific loss of *Itgb8* perturbs MF progression in vivo, we utilized a genetic model by crossing *Itgb8*^fl/fl^ mice with *LepR*^Cre^ mice. We transplanted *LepR*^Cre^*Itgb8*^fl/fl^ and control mice (*LepR*^Cre^*Itgb8*^+/+^ or *Itgb8*^fl/fl^) with c-kit^+^ cells expressing MPL^W515L^ (Fig. 5a). *LepR*^Cre^*Itgb8*^fl/fl^ recipients showed improved BM cellularity, decreased frequency of mutant GFP^+^ cells in BM and PB, and diminished expansion of myeloid cells expressing MPL^W515L^ compared to the control mice (Fig. 5b and Extended Data Fig. 4a-d). In contrast to animals with *Ebf1*-deficient BM niche, the MSC-specific *Itgb8* ablation did not improve splenomegaly (Fig. 5c). Nevertheless, reticulin staining of BM section revealed that *LepR*^Cre^*Itgb8*^fl/fl^ mice exhibited less fibrosis compared to controls, which phenocopies the recipients with *Ebf1*-deficient MSCs (Fig. 5d and Extended Data Fig. 4g). Surprisingly, flow cytometric analysis of BM samples revealed that *LepR*^Cre^*Itgb8*^fl/fl^ recipients retained significantly more hematopoietic progenitors without MPL mutation (LSK-GFP^−^cells; Lineage^−^Sca1^+^c-kit^+^) in their BM, suggesting that reducing fibrosis improves BM niche function and delays the BM marrow failure phase of MF (Extended Data Fig. 4e-f). These results confirm that MSC-specific ITGB8 expression contributes to fibrotic remodeling of the BM niche and MF development.

**Fig. 5.**
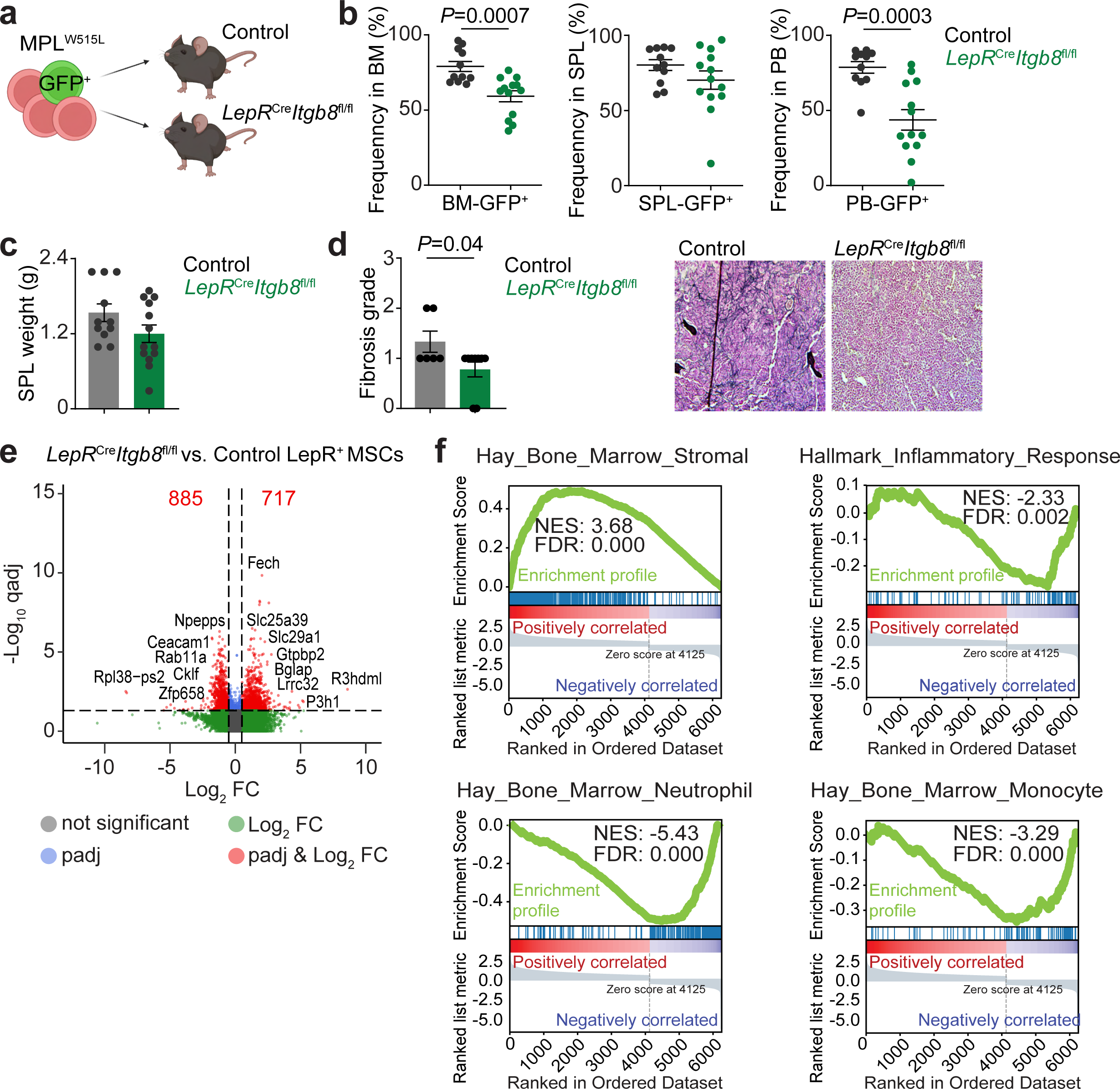
*Itgb8* deficiency in LepR^+^MSCs diminishes MF development. **a.** Experimental schematic. **b.** Frequency of MPL^W515L^ mutant (GFP^+^) cells in BM, SPL, and PB of control and *LepR*^Cre^*Itgb8*^fl/fl^ recipients. **c.** Spleen weights of recipient mice shown in (b). Each dot represents an individual recipient, data represent the mean ± SEM, *n*=11 controls and *n*=13 *LepR*^Cre^*Itgb8*^fl/fl^ mice in 3 independent experiments. **d.** Fibrosis grade scored based on reticulin staining and representative images of reticulin staining in BM sections (40X objective) of control and *LepR*^Cre^*Itgb8*^fl/fl^ mice transplanted with MPL^W515L^ expressing cells. Each dot represents an individual recipient, data represent the mean ± SEM, *n*=6 controls and *n*=9 *LepR*^Cre^*Itgb8*^fl/fl^ mice. **e.** Volcano plot depicting differentially expressed genes in bulk RNA-seq (>2-fold, *P <* 0.05) from LepR^+^ MSCs isolated from *LepR*^Cre^*Itgb8*^fl/fl^ and control mice transplanted with MPL^W515L^ expressing cells (*n* = 3 biologically independent samples per group). The differential gene expression between the conditions was calculated using Cuffdiff. The differentially expressed genes were filtered with the *q* value cutoff <0.05 following Benjamini–Hochberg (FDR, false discovery rate) multiple testing correction of the original *P* values. **f.** Gene set enrichment analysis (GSEA) from *Itgb8*-deficient vs control LepR^+^MSCs purified from *LepR*^Cre^*Itgb8*^fl/fl^ and control mice transplanted with MPL^W515L^ cells. Normalized Enrichment Score (NES), False Discovery Rate (FDR).

To gain insight into molecular alterations underlying the phenotype of *Itgb8*-deficient MSCs, we performed transcriptome analysis of LepR^+^ MSCs isolated from *LepR*^Cre^*Itgb8*^fl/fl^ and control mice two weeks after transplantation with MPL^W515L^ expressing cells. We identified 717 upregulated and 885 downregulated genes (qval≤0.05) in *Itgb8*-deficient MSCs compared to the control MSCs (Fig. 5e). Gene set enrichment analysis (GSEA) and gene ontology (GO) analysis revealed a negative correlation with pathways involved in myeloid leukocyte activation, monocyte and neutrophil development, reflecting reduced frequencies of myeloid cells in the BM of *LepR*^Cre^*Itgb8*^fl/fl^ mice (Fig. 5f and Extended Data Fig. 4h). Moreover, pathway analysis showed positive correlation with signatures for BM stromal cells and ECM organization, implying preserved identity of MSCs and consequently improved BM niche function (Fig. 5f and Extended Data Fig. 4h-i). Additionally, pathways related to inflammatory response, actin cytoskeleton organization, mast cell activation, and production of TNF and TNF superfamily cytokines were enriched in downregulated genes in *Itgb8*-deficient MSCs relative to controls (Fig. 5f and Extended Data Fig. 4h). Interestingly, recent studies revealed that mast cell expansion correlates with the degree of fibrosis in MF mouse models and patients^19, 23, 42^. In MF, mast cells boost the production of TGF-β, TNF, and IL-4/Il-13, leading to increased interactions with megakaryocytes, MSCs, and BM fibroblasts and contributing to BM fibrosis^19, 23, 42^. Hence, these transcriptomic alterations are consistent with the phenotype observed in mice with MSC-specific ITGB8 deletion, where preserving BM niche function ameliorates BM fibrosis, inflammation, and disease burden.

### Pharmacological ITGB8 inhibition halts BM fibrosis and MF progression

To assess whether ITGB8 inhibition might be leveraged therapeutically, we transplanted wild-type C57BL/6J mice with c-kit^+^ cells expressing MPL^W515L^ and subsequently treated them with control IgG or ITGB8 neutralizing antibodies (Fig. 6a). Recipients were monitored for the frequency of GFP^+^ MPL^W515L^ cells in peripheral blood (PB) to ensure equal disease development. Antibodies were administered twice a week (10mg/kg) via intraperitoneal injections starting at 8-10 days post-transplant with GFP^+^ of 8-10% in PB. Similarly to the mice with MSC-specific *Itgb8* deletion, mice treated with ITGB8 neutralizing antibodies exhibited improved BM cellularity and reduced frequency of GFP^+^ MPL^W515L^ cells in the BM, spleen, and PB compared to IgG-treated recipients (Fig. 6b-c and Extended Data Fig. 5a). The reduction in mutant allele burden was associated with diminished expansion of mutant granulocytes (Mac1Gr1^hi^-GFP^+^) and monocytes (Mac1Gr1^lo^-GFP^+^) (Extended Data Fig. 5b-d). ITGB8 inhibition did not affect splenomegaly as spleen weights were comparable between treatment groups, which is also consistent with the genetic model disrupting ITGB8 in MSCs (Extended Data Fig. 5e). Most importantly, recipients treated with ITGB8 neutralizing antibodies showed significantly reduced BM fibrosis, with over 30% of treated mice without any detectable reticulin staining in BM sections (Fig. 6d and Extended Data Fig. 5f). Interestingly, anti-ITGB8 treatment resulted also in decreased levels of multiple inflammatory mediators in the BM lavage such as TGF-β, IL-1a, IL-b, S100A8, S100A9, IL-4, IL-6, TNFα, CXCL1, TIMP-1 and IL-12p17, which are normally elevated in MF mouse models and patient samples (Fig. 6e and Extended Data Fig. 5g). Systemic inflammation is a known hallmark of MF, suggested to drive disease progression. Many of these inflammatory factors are known to promote tissue fibrosis via direct or indirect activation of fibroblasts, and targeting these cytokines has been reported to elicit therapeutic benefits in MF mouse models and patients^8, 23–25, 43–45^. Moreover, the levels of myeloid growth factors G-CSF and M-CSF were also reduced by blocking ITGB8, which correlates with decreased expansion of granulocytes and monocytes in treated recipients (Fig. 6e and Extended Data Fig. 5b-c). Considering that most of these factors are produced by damaged MSCs and fibroblasts during MF, decreased BM inflammation along with reduced fibrosis indicates a markedly improved BM niche function in MF recipients treated with ITGB8 neutralizing antibodies. Furthermore, anti-ITGB8 treatment resulted in increased frequency and absolute numbers of non-mutant hematopoietic progenitors (LSK-GFP^−^ cells) relative to controls (Extended Data Fig. 5h-i), demonstrating improved BM niche function capable of sustaining normal hematopoiesis and again recapitulating the genetic model. This is in line with recent reports suggesting that anti-tumor effects of TGFβ inhibition might be partially achieved via reactivation of normal hematopoiesis, whereby healthy cells can compete with mutant ones^46^. Given that the BM niche is crucial to maintain healthy HSCs, our results emphasize the importance of preserving its proper function in MF and other hematological disorders.

**Fig. 6.**
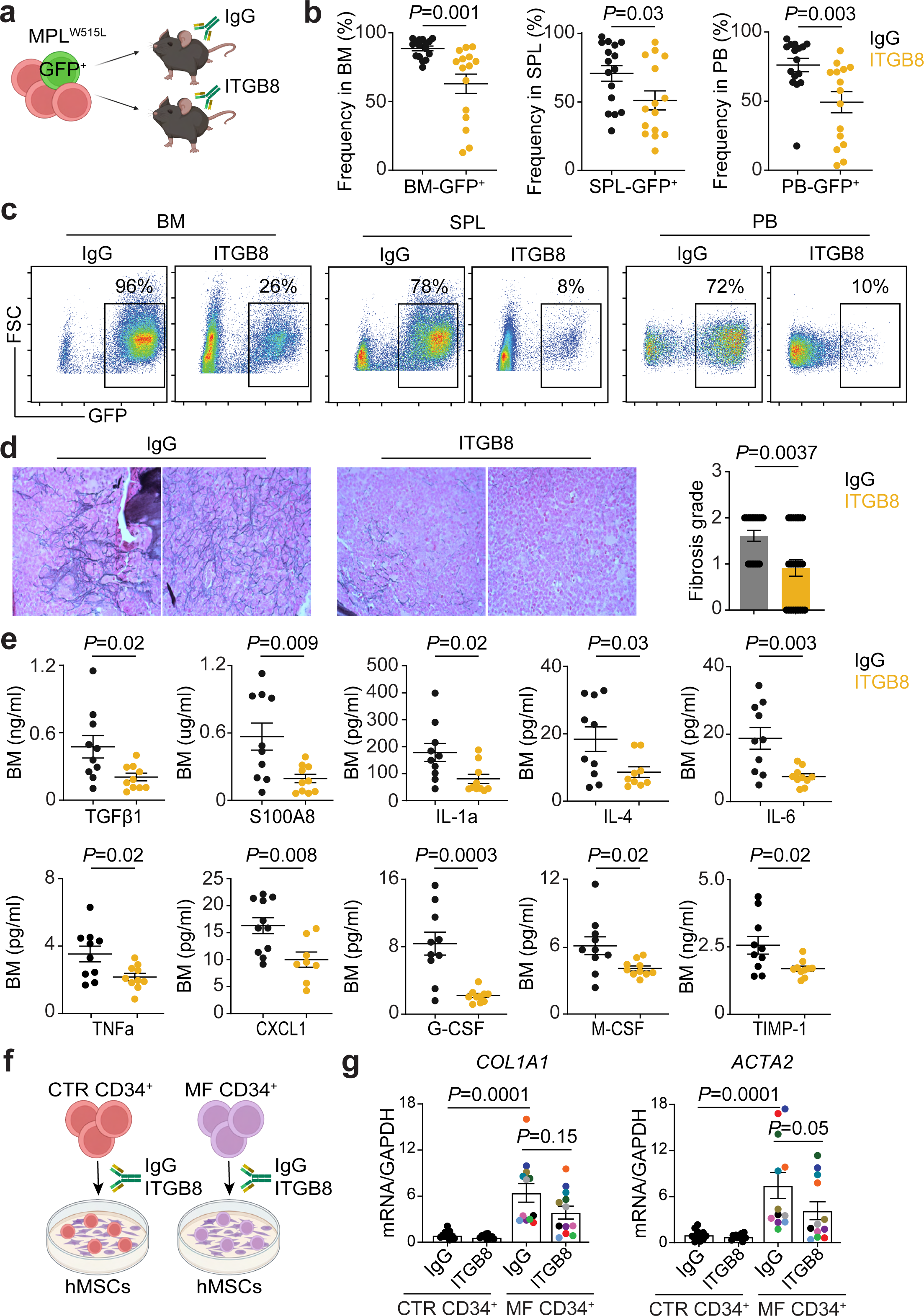
Pharmacological ITGB8 inhibition reduces MF progression. **a.** Experimental setup. **b.** Frequencies of GFP^+^ mutant cells (MPL^W515L^) in BM, SPL, and PB of mice treated with control IgG or ITGB8 neutralizing antibodies (10mg/kg). Each dot represents an individual recipient, data represent the mean ± SEM, *n*=16 IgG treated and *n*=15 anti-ITGB8 treated mice in 3 independent experiments. **c.** Representative flow cytometry plots for data shown in (b). **d.** Fibrosis grade scored based on reticulin staining and representative images of reticulin staining in BM sections (40X objective) of mice transplanted with MPL^W515L^ expressing cells and treated with IgG or ITGB8 neutralizing antibodies. Each dot represents an individual mouse, data represent the mean ± SEM, *n*=18 IgG treated and *n*=23 anti-ITGB8 treated mice. **e.** Cytokine levels quantified by multiplex cytokine array (Luminex) in the BM supernatants isolated from mice transplanted with MPL^W515L^ cells and treated with control IgG or ITGB8 neutralizing antibodies. Each dot represents an individual recipient, data represent the mean ± SEM, *n*=10 IgG treated and *n*=8-10 anti-ITGB8 treated mice. **f.** Experimental setup. **g.** Relative mRNA levels of fibrotic markers in human MSCs co-cultured for 72h with CD34^+^ progenitors from MF patients (MF CD34^+^) or age- and gender-matched healthy donors (CTR CD34^+^) in the presence of IgG or anti-ITGB8 antibodies (10µg/ml). hMSCs were derived from BM of 6 independent healthy donors. CD34^+^ cells were enriched from MF patients harboring mutations in *JAK2* (3 patients), *MPL* (5 patients), and *CALR* (4 patients). Each dot represents an individual donor, and each MF patient is represented by different color dot. Data represent the mean ± SEM, *n*=11 CTR CD34^+^ and *n*=12 MF CD34^+^ donors in 4 independent experiments.

To evaluate whether blocking of ITGB8 elicits beneficial effects on human cells, we leveraged co-cultures of hMSCs with CD34^+^ cells from healthy donors (CTR CD34^+^) or MF patients (MF CD34^+^) in the presence of control IgG or ITGB8 neutralizing antibodies (Fig. 6f). In the control IgG group, expression of hallmark fibrotic markers *COLA1A1* and *ACTA2* was increased in hMSC co-cultured with MF CD34^+^ cells relative to co-cultures with healthy CD34^+^ cells. However, upon ITGB8 inhibition, expression of both fibrotic genes was decreased in hMSCs co-cultured with MF CD34^+^ cells (Fig. 6g). CD34^+^ cells used for co-culture experiments were derived from MF patients harboring all three driver mutations (*JAK2*, *MPL*, or *CALR*), indicating that mechanisms underlying fibrotic remodeling are mutation-agnostic. Collectively, these results show that ITGB8 has therapeutic potential to alleviate BM fibrosis and halt MF progression.

### Hematopoietic-specific Itgb8 deletion does not affect MF development

*Itgb8* is also expressed by the hematopoietic compartment, especially T cells, where it mediates tumor immune escape via TGFβ activation^27, 28^. ITGB8 inhibition, along with immune checkpoint inhibitors, has been successfully applied as immunotherapy in preclinical models of multiple solid tumors^28^. Moreover, blocking TGFβ activation via regulatory T cells has been reported to elicit anti-tumor activity in MPNs^46^. Therefore, we also examined whether hematopoietic-specific *Itgb8* deletion affects MF development. Hematopoietic progenitors (c-kit^+^ cells) isolated from *Vav1*^Cre^*Itgb8*^fl/fl^ and control mice (*Vav1*^Cre^*Itgb8*^+/+^ or *Vav1*^fl/fl^) were transduced with the MPL^W515L^ construct and transplanted into wild-type (CD45.1) recipients (Extended Data Fig.6a). Mice that received *Itgb8*-deficient hematopoietic cells harboring the MPL mutation did not show any differences in disease burden, spleen weights or BM fibrosis compared to the controls (Extended Data Fig. 6b-e). These data indicate that ITGB8 inhibition in blood cells is not sufficient to modulate MF.

## Discussion

Bidirectional communication between hematopoietic and BM niche cells is crucial for BM homeostasis. In malignancies, however, neoplastic blood cells modify healthy MSCs via direct cell–cell contact and secreted factors, creating a dysfunctional hematopoietic microenvironment^7, 11, 47^. Alterations in the BM niche have been described in acute lymphoblastic leukemia (ALL), acute myeloid leukemia (AML), myelodysplastic syndrome (MDS), and MPNs^7, 47^. The corrupted BM microenvironment confers a competitive advantage to malignant HSCs over healthy ones, facilitating immune evasion and chemoresistance. Our work and the existing literature support the premise that targeting the aberrant crosstalk between malignant cells and their microenvironment, in combination with current standards of care, may provide a promising strategy for treating hematological malignancies.

Although MF originates within hematopoietic cells, progressive BM fibrosis has been recognized as a critical characteristic contributing to MF pathogenesis. Initiated by blood cells harboring *JAK2*, *MPL* or *CALR* mutations, fibrotic remodeling of the BM niche renders a hostile environment for normal hematopoiesis, leading to a variety of clinical symptoms and poor outcomes for patients^12, 13^. Current pharmacological approaches, while helpful with constitutional symptoms, do not improve BM fibrosis or eradicate the mutant clones^16, 17^. The reported rates of cessation of therapy with ruxolitinib, a JAK2 inhibitor, among MF patients are 49%, 71%, and 86% after year 1-3, respectively, and the median overall survival after treatment discontinuation is 11-16 months^18, 48^. Thus, there is an urgent need to identify new therapeutic targets and design more effective treatments that will address not only the mutant initiating clones but also the defective BM microenvironment. Recent studies revealed that bone marrow LepR^+^ MSCs undergo fibrotic conversion during MF, shedding light on the origin of the myofibroblasts that drive fibrosis^8, 9, 19^. These studies advanced our understanding of the hematopoietic cell types and signals they send to communicate with the BM niche, which drives BM fibrosis. Nevertheless, the MSC-specific factors that respond to these hematopoietic-derived signals and subsequently control transcriptional response resulting in fibrotic remodeling remain understudied.

In this report, we identified transcription factor EBF1 as a master regulator of the fibrotic gene program in BM MSCs. We show that EBF1 expression is increased in both human and mouse MSCs in response to hematopoietic cells carrying each of the three MPN driver mutations. Moreover, intersecting the EBF1 chromatin occupancy and transcriptome analysis revealed that EBF1 binds regulatory elements of genes responsible for fibrosis, indicating that it is a direct transcriptional regulator of fibrosis. Furthermore, MSC-specific deletion of EBF1 protected mice from developing BM fibrosis and lead to a reduced disease burden in mice transplanted with MPL^W515L^-expressing hematopoietic progenitors. Interestingly, a recent study profiling solitary fibrous tumor—mesenchymal-derived soft tissue tumors characterized by overproduction and pathological remodeling of ECM—reported hypomethylation of the EBF1 locus and increased expression of EBF1 in these tumors^49^. Also, a study investigating fibroblast heterogeneity in pulmonary fibrosis uncovered an EBF1-positive fibroblast population that substantially contributes to ECM deposition in multiple models of lung fibrosis^50^. These observations support our proposal of EBF1 as a master regulator of fibrotic remodeling and suggest that the EBF1-controlled transcriptional program we pinpointed in BM MSCs is conserved in stromal and fibroblast-like cells across different tissues.

Importantly, we identified ITGB8 as an EBF1-regulated gene with potential to serve as target in MF directed therapies. We show that ITGB8 inhibition with neutralizing antibodies results in less BM fibrosis, reduced expansion of MPL-mutant cells, and decreased BM inflammation. Using genetic models, we also verified that ITGB8 inhibition in BM MSCs halt MF progression. This result reinforces the hypothesis that preventing or reducing BM fibrosis might be beneficial in ameliorating MF. It is worth mentioning that ITGB8 inhibition in the tumor microenvironment has been successfully used to overcome TGF-β-induced immunosuppression in preclinical mouse models of multiple solid tumors^28^. Moreover, there are currently several active clinical trials evaluating the utility of ITGB8 blockade for the treatment of solid tumors.

Our data indicate that inhibiting the EBF1-ITGB8 axis in MF MSCs in combination with other treatment options that target mutant blood cells might constitute a novel therapeutic approach. Furthermore, reducing fibrosis and improving BM microenvironment function might favor the engraftment of allogeneic donor grafts in MF patients undergoing transplantation. MF patients (intermediate-2 and higher risk) treated with allogeneic stem cell transplantation have a long-term survival advantage, but at the cost of early transplant-related mortality, mostly due to graft failure or delayed engraftment^51^. It is plausible that improving MSC and BM niche function in MF patient transplant recipient might enhance HSC engraftment^52^. Also, given that our data suggest that fibrotic remodeling regulated via the EBF1-ITGB8 axis is mutation-agnostic, it is possible that this pathway could be exploited to treat other blood disorders such as MDS with BM fibrosis. MDS patients with moderate-to-severe fibrosis show more profound multilineage cytopenia, worse response to azacitidine, faster leukemic progression, and worse survival compared with patients with no or mild fibrosis^53, 54^. Similarly to MF, MDS patients with fibrosis that undergo allogeneic HSC transplantation present with delayed engraftment, higher rates of relapse, and lower event-free survival compared to the MDS transplant recipients without BM fibrosis^55–57^. Currently, there are no approved treatments available to reverse BM fibrosis in MF patients other than allogeneic stem cell transplantation. However, there are ongoing clinical trials targeting collagen crosslinking in MF patients (ClinicalTrials.gov Identifier: NCT04676529, NCT04679870) aimed at improving the BM microenvironment in MF patients. Our study contributes to such efforts and warrants further investigation using ITGB8 inhibition as an anti-fibrotic treatment in MF patients.

## Acknowledgments

We are grateful to Sandra Capellera-Garcia and Eirini Trompouki for reading the manuscript and critical comments. We also thank members of Division of Experimental Hematology at St. Jude Children’s Research Hospital for valuable feedback on the study. We thank institutional core facilities (Animal Resource Center, Comparative Pathology Core, Flow Cytometry and Cell Sorting, Hartwell Center for Sequencing, and Biostatistics) without whom this work would not be possible. In particular, we thank Dr. Peter Vogel for his expertise in pathology and fibrosis scoring. We also thank David Cullins, Emilia Kooienga, and Hematology Flow Core Facility for technical assistance and the discussions regarding experimental design. We are also grateful to Anna Baird, Shelby Patrick, Clemothy Bell, Paisley Creedon, MyEsha Bingham, and Kristen Philippart for their help in Animal Resource Center. This work was supported by the following grants: Leukemia Research Foundation (MD), When Everyone Survives Foundation (MD), P30 CA021765 (National Cancer Institute) (MD), P01 CA108671 (National Cancer Institute) (J.D.C, B.M, R.H), American Lebanese Syrian Associated Charities (ALSAC) (MD, J.D.C).

## Author Contributions

L.T and M.D designed, performed, and analyzed experiments. N.M supported experimental work. R.M performed bioinformatics analysis. L.K and T.L performed CUT&RUN experiments. B.M and R.H provided PBMCs from MF patients. D.S and A.A generated, characterized and provided ITGB8 neutralizing antibodies. P.V performed pathology analysis. J.D.C supervised L.K and T.L. M.D conceived and supervised the study. M.D wrote the manuscript with input from all authors.

## Methods

### Mice

*Ebf1*^fl/fl^ mice were generated and provided by Dr. R. Grosschedl (Max Planck Institute, Freiburg, Germany)^34^. *Itgb8*^fl/fl^ mice, (Stock *Itgb8^tm2Lfr^*/Mmucd, RRID:MMRRC_014108-UCD) were obtained from the Mutant Mouse Resource and Research Center (MMRRC) at University of California at Davis, an NIH-funded strain repository, and was donated to the MMRRC by Dr. Louis Reichardt (University of California, San Francisco)^58^. *Prx1*^Cre^ (Strain #005584), *LepR*^Cre^ (Strain #008320), *Vav1*^Cre^ (Strain #008610), CD45.1-C57BL/6J-Ptprcem6Lutzy/J (JAXBoy, Strain #033076), and wild-type C57BL/6J (Strain #000664) mice were purchased from Jackson Laboratory. Unless indicated otherwise, 8-14 week-old male and female mice were used for experiments. All mice were bred and housed under specific pathogen-free (SPF) conditions. All animal procedures were performed according to protocols approved by the St. Jude Children’s Research Hospital Institutional Animal Care and Use Committee.

### Retrovirus production

Retroviral constructs MSCV-MPL-WT-IRES-GFP and MSCV-MPL-W515L-IRES-GFP were generously provided by Dr. J. Crispino^32^. Retroviruses were produced in Plat-E Cells (Cell Biolabs) by transfection with 10 μg of plasmid using FuGENE HD transfection reagent (Promega) in Opti-MEM medium (Gibco). After overnight incubation, the medium was replaced with RPMI containing 5% fetal bovine serum (FBS). The virus supernatant was collected at 48h post transfection.

### Retroviral transduction

Bone marrow cells were isolated from JAXBoy or C57BL/6J mice by crushing the bones (tibias, femurs, hip bones, and spines) in FACS buffer (PBS-2% FBS) using pestle and mortar. C-kit positive hematopoietic progenitors were enriched using CD117 magnetic microbeads and autoMACS separator (Miltenyi Biotec). Cells were cultured in StemSpan media (Stem Cell Technologies) with murine SCF (50ng/ml), human IL-6 (10ng/ml), murine IL-3 (10ng/ml), human LDS (1:500 dilution), and penicillin/streptomycin. After an overnight culture at 37°C, cells were transduced with MPL^WT^ or MPL^W515L^ retrovirus supernatants supplemented with murine SCF (50ng/ml), human IL-6 (10ng/ml), murine IL-3 (10ng/ml), human LDL (1:500) and LentiBOOST transduction enhancer (1:100 dilution). Cells were spinoculated with viral supernatants at 2,500 RPM for 1.5 hours at 32°C and then incubated with the viral supernatant for 3 hours at 37°C. Next, the media was changed to StemSpan (Stem Cell Technologies) with murine SCF (50ng/ml), human IL-6 (10ng/ml), murine IL-3 (10ng/ml), human LDL (1:500 dilution), and penicillin/streptomycin, and cells were incubated overnight at 37°C. The transduction efficiency was based on the percentage of GFP positive cells (GFP^+^) measured by flow cytometry.

### Transplantation experiments

Recipient mice (8-14 week-old) were sub-lethally irradiated with 5.5 Gy using a gamma irradiator with a ^137^Cs source. Recipients were treated with Sulfamethoxazole/Trimethoprim in drinking water for 7 days before the transplant and for 21 days post-transplant. For transplantation of c-kit^+^ cells expressing MPL^W515L^ into wild-type recipients or mice crossed to *Prx1*^Cre^ and *LepR*^Cre^ mice, c-kit^+^ cells were isolated from CD45.1 JAXBoy mice. For transplant experiments using *Vav1*^Cre^ mice, c-kit^+^ cells were isolated from *Vav1*^Cre^*Itgb8*^fl/fl^ and control mice (*Vav1*^Cre^*Itgb8*^+/+^ or *Vav1*^fl/fl^) and transduced with MPL^W515L^ retrovirus and then transplanted into CD45.1 JAXBoy recipients. 3×10^5^ of MPL^W515L^ GFP+ cells were injected into the tail vein of recipient mice. Unless otherwise specified, peripheral blood was collected two weeks post-transplant via retro-orbital sampling for flow cytometry analysis. Mice were euthanized 4-5 weeks post-transplant.

### ITGB8 neutralizing antibody treatment

Wild type CD45.1 JAXBoy mice were transplanted with c-kit^+^ cells expressing MPL^W515L^ as described above. Peripheral blood was collected on day eight post-transplant and analyzed by flow cytometry to estimate the engraftment of GFP+ MPL^W515L^ cells. Animals were randomized into treatment groups. The control mice received the IgG antibodies (10 mg/kg), and the experimental mice were injected with ITGβ8 neutralizing antibodies (10 mg/kg) via intraperitoneal injections twice a week for 3 weeks starting on day 9 post transplantation. ITGβ8 neutralizing antibodies (mouse monoclonal) were provided by Dr. Dean Sheppard (UCSF)^28^. Peripheral blood was collected via retro-orbital sampling for flow cytometry and CBC analysis every week to monitor disease progression based on GFP frequency. Mice were sacrificed 40 days post-transplant. Peripheral blood, bone marrow, and spleen were collected for flow cytometry analysis.

### Cell culture and RNA interference

Intact bones (tibia, femur and hip bones) were isolated and crushed in small fragments in FACS buffer (PBS+2%FBS) using a mortar and pestle. BM cells were collected and red blood cells lysis was performed (#420302, Biolegend). Cells were washed with FACS buffer and CD45 depletion was performed using magnetic MojoSort Mouse CD45 Nanobeads (#480028, Biolegend) according to manufacturer’s instruction. Bone chips resulting from the crushing were digested using 0.4% collagenase type II (#LS004176, Worthington) + 0.01% DNase I (# 04536282001, Roche) in DMEM for 45min at 37°C with gentle shaking. After digestion, the cells were filtered through 70 µm strainer, washed, and depleted from CD45+ cells using magnetic beads as described before. Combined BM cells and cells from bone chips digestion were stained for FACS-sorting. FACS-sorted LepR^+^MSCs were cultured at 37 °C and 3% O_2,_ 5% CO_2_ in MesenCult™ Expansion Kit media (Stem Cell Technologies) supplemented with L-Glutamine and penicillin/streptomycin. Human MSCs (hMSCs) were purchased from Lonza and cultured at 37 °C and 3% O_2,_ 5% CO_2_ in MesenCult™ Proliferation Kit media (Stem Cell Technologies) supplemented with penicillin/streptomycin. Sub-confluent MSCs were transfected with 50 nM of ON-TARGETplus SMARTpool siRNA (Dharmacon, Horizon Discovery) against mouse *Ebf1* (L-045017-01), mouse *Itgb8* (L-066709-01), human *EBF1* (L-011848-01), human *ITGB8* (L-008014-00) or Non-targeting Control Pool (D-001810-10), according to the manufacturer’s instructions. Transfections were carried out in respective cell culture media without antibiotics, using DharmaFECT 1 Transfection reagent.

### Isolation of CD34^+^ cells from patients’ peripheral blood

Frozen vials of PB from MF patient or healthy controls were thawed in a water bath at 37°C and washed with 20ml of StemSpan medium (#02691, StemCell). Cells were centrifuged at 400g for 5min and CD34^+^ cells isolation was performed using the EasySep™ Human CD34 Positive Selection kit (#17856, StemCell) according to manufacturer’s instructions. Peripheral blood mononuclear cells from MF patients were provided by Drs. Bridget Marcellino and Ronald Hoffman and MNPR Consortium at Icahn School of Medicine at Mount Sinai, New York. The age-and gender-matched peripheral blood mononuclear cells from healthy donors were purchased from Stem Cell Technologies.

### Co-culture experiments

FACS-sorted LepR^+^MSCs were expanded for 12-14 days at 37 °C and 3% O_2,_ 5% CO_2_ in MesenCult™ Expansion Kit media (Stem Cell Technologies) supplemented with L-Glutamine and penicillin/streptomycin. 2×10^5^ of LepR^+^MSCs were seeded in 6-well plates and transfected with siRNAs the next day. 2×10^5^ of MPL^WT^ or MPL^W515L^ expressing c-kit^+^ cells were plated on LepR^+^MSCs 48 hours after siRNA-mediated knockdown of *Itgb8* and co-cultures were carried out for 72 hours at 37 °C and 3% O_2,_ 5% CO_2_ in StemSpan (Stem Cell Technologies) supplemented with murine SCF (50ng/ml), human IL-6 (10ng/ml), murine IL-3 (10ng/ml), and penicillin/streptomycin. After 72h, LepR^+^MSCs were collected by incubation with trypsin-EDTA and CD45+ cells were depleted using MojoSort Mouse CD45 nanobeads (BioLegend). RNA was isolated and used for qPCR.

hMSCs were expanded for 7-10 days at 37 °C and 3% O_2,_ 5% CO_2_ in MesenCult™ Proliferation Kit media (Stem Cell Technologies) supplemented with penicillin/streptomycin. 8×10^4^ of hMSCs were seeded in 12-well plates coated with MesenCult™-SF Attachment substrate (Stem Cell Technologies) and transfected with siRNAs the next day. 2×10^5^ of CD34^+^ cells purified from peripheral blood of healthy donors of MF patients were plated on hMSCs 48 hours after siRNA-mediated knockdown of *EBF1* and co-cultures were carried out for 72 hours at 37 °C and 3% O_2,_ 5% CO_2_ in StemSpan with CD34+ Expansion Supplement 10X (Stem Cell Technologies) and penicillin/streptomycin. After 72h, hMSCs were collected by incubation with trypsin-EDTA and CD45+ cells were depleted using MojoSort Human CD45 nanobeads (BioLegend). RNA was isolated and used for qPCR. Co-cultures with ITGB8 neutralizing antibodies were carried out in the same conditions in the presence of control IgGs (10µg/ml) or anti-ITGB8 antibodies (10µg/ml).

### Protein extraction and Western blot

Cell pellets were lysed in RIPA buffer (#89901, Thermofisher) supplemented with protease/phosphatase inhibitors (78440, Thermofisher) for 20min at 4°C with agitation. Lysates were centrifuged at 16000g for 10min at 4°C and supernatants were collected, aliquoted and stored at -80°C. Protein concentration was measured using Pierce™ BCA Protein Assay Kit (#23225, Thermofisher). Samples containing 15-30 ug of protein were separated on a 4-20% SDS-PAGE gel under denaturing conditions and transferred on a PVDF membrane using Trans-Blot® SD Semi-Dry Electrophoretic Transfer Cell. The blotted membranes were blocked with TBST+5% BSA for 2h at RT on an orbital shaker, followed by incubation with respective primary antibodies against EBF1 (1/1000, rabbit polyclonal)^34^ and ITGB8 (1/500, GTX64493 Genetex) in TBST+3% BSA at 4°C overnight on an orbital shaker. Membranes were then washed three times with TBST buffer for 10 min each on an orbital shaker and probed with the secondary antibody anti-rabbit IgG-HRP (1:2000) and signals were visualized using SuperSignal™ West Pico PLUS Chemiluminescent Substrate (#34580, Thermofisher). Membranes were next washed extensively with TBST buffer and reprobed with anti-beta Actin-HRP (1/10000, ab49900 Abcam).

### Flow cytometry and cell sorting

Intact bones (tibias, femurs, hip bones) were cut into small fragments and crushed in FACS buffer (PBS-2% FCS) using pestle and mortar. Bone marrow cells were collected and red blood cell lysis was performed. Next, the cells were washed with FACS buffer and stained for analysis. Spleens were crushed in FACS buffer through the 70 µm mesh filter with the cell strainer pestle. Cells suspension was filtered through 70 µm filter twice to achieve single cell suspension. After red blood cell lysis, cells were washed with FACS buffer and stained for flow cytometry analysis. Peripheral blood was collected in tube pretreated with K2EDTA (BD Microtainer; Becton, Dickinson and Co, 365974) and treated with the red blood cell lysis buffer prior to washing with the FACS buffer and staining for analysis. Data were acquired using LSR Fortessa or FACSymphony™ A3 (BD Biosciences). Data analysis was performed using FloJo Version 10.8 (LLC). BD FACSAria Fusion and BD Symphony S6 were used for FACS-activated cell sorting. Supplementary Table 1 contains a full list of used antibodies.

### Histopathology

Tissues were fixed in 10% neutral buffered formalin (Globe Scientific) and processed at the Comparative Pathology Core at St. Jude Children’s Research Hospital. Reticulin staining was scored blindly.

### Cytokine analysis

BM fluids were collected from Intact bones (tibia, femur and hip bones) isolated from mice treated with IgG or ITGB8 neutralizing antibodies and crushed in PBS (# 10010-023, Gibco) using a mortar and pestle. Cells were centrifuged at 400g for 5 min at 4°C. The supernatants were transferred to the new tubes, centrifuged at 8000g for 10 min at 4°C, transferred again to fresh tubes and stored at -80°C. The cytokine analysis was performed by Eve Technologies using Luminex 200 system.

### RNA extraction and real-time qPCR

Total RNA was isolated with Quick-RNA Microprep Kit (Zymo Research) and treated with DNase (Qiagen) according to the manufacturer’s instructions. Complementary DNA (cDNA) was synthesized with the SuperScript II Reverse Transcriptase Kit (Invitrogen) and real-time qPCR was performed with the Fast SYBR Green Master Mix (Applied Biosystems) according to manufacturer’s guidelines. All the samples were assayed in duplicates and analyzed with a Step One Plus Real-Time PCR System (Applied Biosystems). Supplementary Table 1 contains a full list of the primer sequences.

### Bulk RNA-seq and data analysis

Total RNA was prepared from FACS-sorted LepR^+^MSCs using Arcturus PicoPure RNA Isolation Kit (Applied Biosystems) according to manufacturer’s instructions. Three biological replicates were used for RNA-seq from each genotype and 5-6×10^3^ of LepR^+^MSCs were used per replicate. Library preparation and sequencing were performed by Vanderbilt Technologies for Advanced Genomics Core. RNA quality was evaluated using an Agilent Bioanalyzer. The SMART-seq v4 Ultra Low Input RNA kit was used to prepare sequencing libraries and the NovaSeq X Plus system was used to generate paired-end, 150 bp reads at the depth of 50 M reads per sample. Bulk RNA-seq data were processed using a custom pipeline (documented at HEMTools), utilizing Kallisto (v0.43.1) for pseudo-alignment to the mm10 reference genome and transcript quantification, and Sleuth (v0.30.0) for differential gene expression analysis. For Gene Set Enrichment Analysis (GSEA), gseapy (V.1.1.8) was used to find the enriched pathways.

### CUT&RUN

Intact bones (tibia, femur and hip bones) were isolated from wild-type C57BL/6J mice and crushed in small fragments in FACS buffer (PBS+2%FBS) using a mortar and pestle. BM cells were collected and red blood cells lysis was performed (#420302, Biolegend). Cells were washed with FACS buffer and CD45 depletion was performed using magnetic MojoSort Mouse CD45 Nanobeads (#480028, Biolegend) according to manufacturer’s instruction. Bone chips resulting from the crushing were digested using 0.4% collagenase type II (#LS004176, Worthington) with 0.01% DNase I (# 04536282001, Roche) in DMEM for 45min at 37°C with gentle shaking. After digestion, the cells were filtered through 70 µm strainer, washed, and depleted from CD45+ cells using magnetic beads as described before. Combined BM cells and cells from bone chips digestion were stained with biotinylated CD11b and CD11c antibodies for 30 min at 4°C and washed with FACS buffer. Next, the cells were incubated with MojoSort streptavidin nanobeads (# 480016, BioLegend) according to manufacturer’s instruction and CD11b/CD11c positive cells were depleted using magnetic rack. The remaining MSCs were expanded at 37 °C and 3% O_2,_ 5% CO_2_ in MesenCult™ Expansion Kit media (Stem Cell Technologies) supplemented with L-Glutamine and penicillin/streptomycin for three passages.

CUT&RUN was performed on MSC nuclei using the CUTANA protocol v1.5(EpiCypher 14-1048). Nuclei were prepared from three biological replicates (500,000 cells per replicate) by incubating mouse mesenchymal cells for 10 minutes in fresh nuclei extraction (NE) buffer consisting of 235 µL pre-nuclei extraction Buffer (# 21-1026, EpiCypher), 0.13 µL 1 M Spermidine, 9.8 µL 25X protease inhibitor cocktail and 24 µL of 10X Phostop per sample. Nuclei were pelleted by centrifugation at 600g for 3 minutes at 4°C and were resuspended in 100 µL NE buffer per sample. For each CUT&RUN reaction,100 µL nuclei were bound to ConA magnetic beads and incubated overnight at 4°C with gentle rocking in the presence of IgG antibody (1:100) and EBF1 rabbit polyclonal antibody (1:50). After washing to remove unbound antibody, pAG-MNase (Protein A/G-Micrococcal Nuclease fusion protein) was added. Targeted chromatin digestion was initiated by the addition of 100 mM calcium chloride and incubated on ice for 2 hours at 4°C. The reaction was stopped by adding STOP Buffer. DNA digested chromatin fragments were released into the supernatant by incubating the tubes at 37°C and subsequently purified using SPRIselect beads (Beckman Coulter). The CUT&RUN derived DNA was quantified using Qubit 1X dsDNA HS Assay and 5 ng of DNA was used to prepare sequencing libraries with CUT&RUN library Prep Kit (# 14-1001, EpiCypher). DNA fragments were first end-repaired and A-tailed, followed by ligation to Illumina-compatible, indexed adapters. DNA was purified using SPRIselect beads and eluted in 15 µL of 0.1X TE buffer and used for indexing PCR. The amplified libraries were purified again with SPRIselect beads and assessed for size distribution (∼300 bp) using an Agilent TapeStation. The final libraries were sequenced on Novaseq Illumina platform using paired-end 50 bp reads. Spike-in DNA and control IgG samples were used to assess digestion specificity and enable normalization across samples.

### CUT&RUN analysis

Community driven nf-core/cut&run pipeline (V.3.1) with nextflow (v.23.04.1) was utilized for CUT&RUN analysis. In detail, pair-end reads were aligned to mm10 reference genome using bowtie2 (V.2.4.4). MACS2(V. 2.2.7.1) was used for peak calling and bamCoverage function from deeptools(V. 3.5.1) was used to generate bigWig files. findMotifsGenome.pl command from homer (V. v5.1) were used for the motif analysis, whereas annotatePeaks.pl command was used for peak annotation.

### Analysis of published scRNA-seq datasets

Public scRNA-seq data were obtained from sources listed below and where reanalyzed as described in the original articles. Tikhonova et al., (PMID: 30971824), GEO accession number GSE108892. Leimkühler et al., (PMID: 33301706), GEO accession number GSE156644. Li, Colombo, Wang et al., (PMID: 39383242), GEO accession number GSE228995.

### Statistical analysis

Statistical analysis was performed using two-tailed unpaired Student’s *t*-test (confidence interval of 95%) with significance of * *P* <0.05, ** *P* <0.01, *** *P* <0.001. Pearson correlation coefficients were calculated using a confidence interval of 95%, two-tailed. A Welch two sample t-test and Wilcoxon rank sum exact test were used to calculate the expression of fibrotic markers in Fig. 6g.

### Data availability

All genome-wide data in this study were deposited at the Gene Expression Omnibus (GEO) under series GSE301409.

---

**Extended Data Fig. 1.**
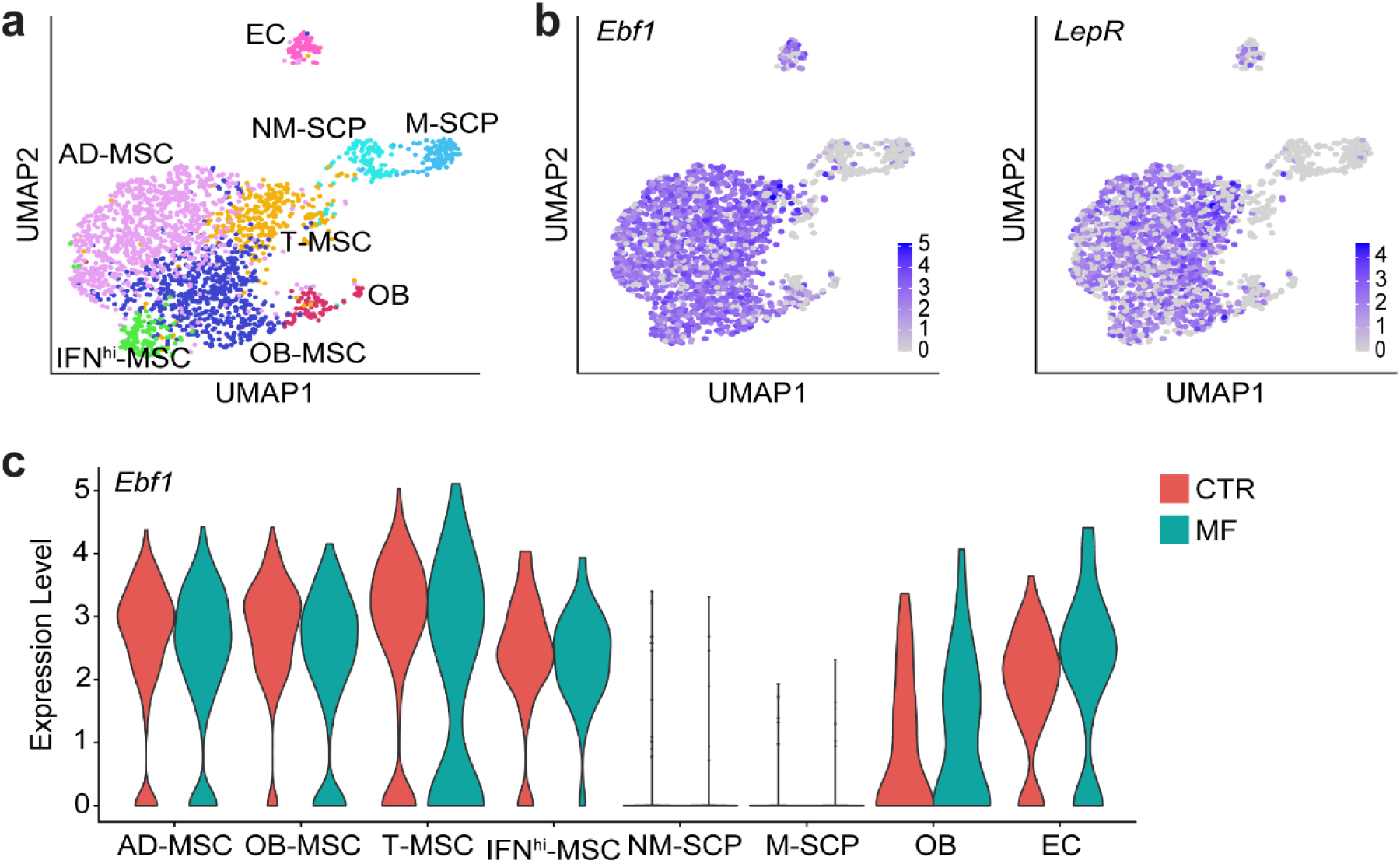
Increased *Ebf1* expression in MPN stroma. **a.** UMAP of non-hematopoietic BM niche cells isolated from wild-type mice transplanted with hematopoietic progenitors expressing GFP (CTR) or thrombopoietin (MF) colored by annotated cell cluster obtained by unsupervised clustering, as reported by Leimkuhlet et al.; adipogenic mesenchymal stromal cells (AD-MSCs); osteogenic mesenchymal stromal cells (OB-MSCs); transitioning mesenchymal stromal cells (T-MSCs); interferon-responsive mesenchymal stromal cells (IFN^hi^-MSCs); non-myelinating Schwann sell progenitors (NM-SCP); myelinating Schwann sell progenitors (M-SCP); osteoblastic cells (OB); endothelial cells (EC). **b.** UMAP of normalized *Ebf1* and *LepR* expression in MSC clusters shown in (a). **c.** Violin plot showing normalized *Ebf1* expression in cell clusters identified in (a).

**Extended Data Fig. 2.**
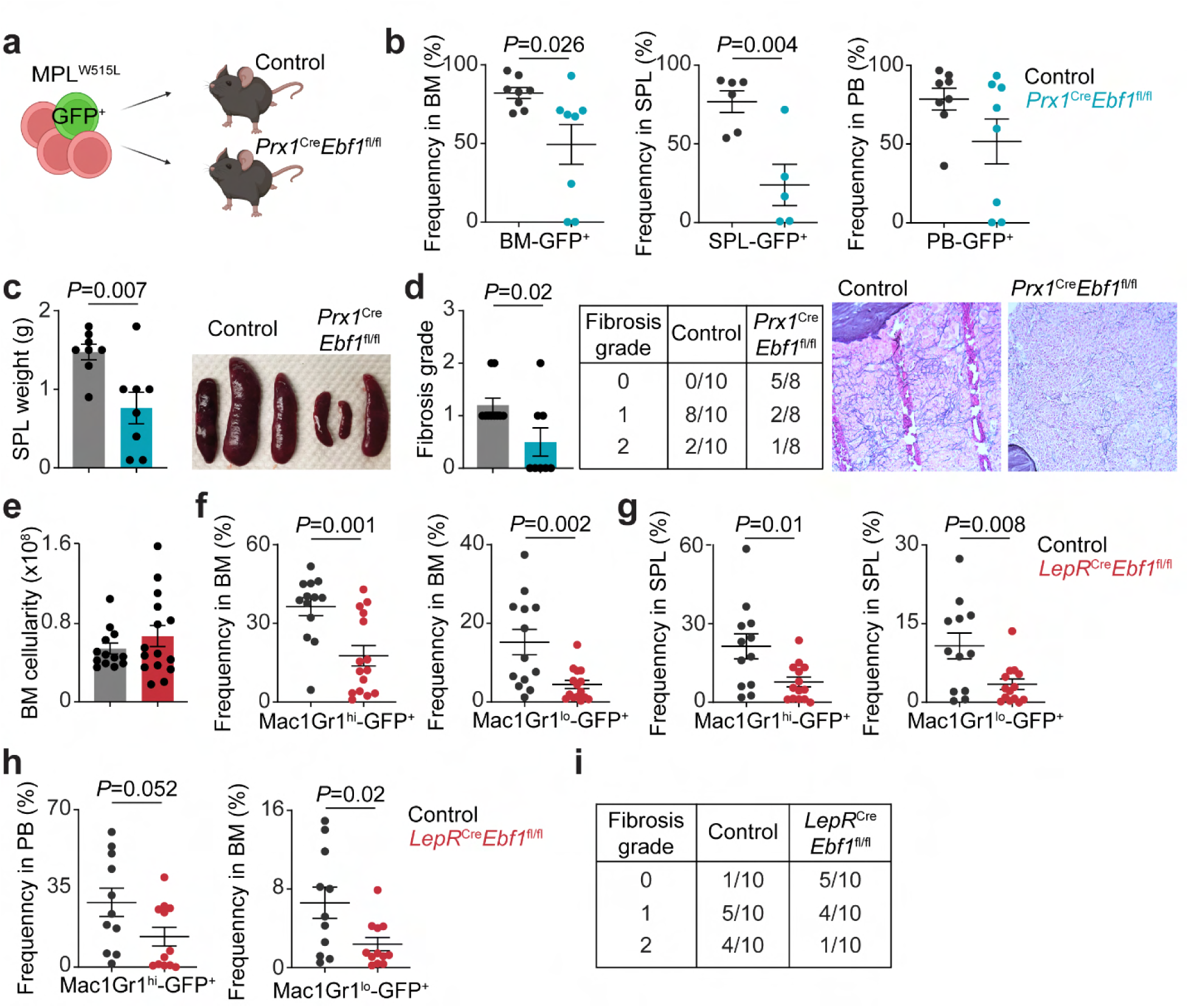
*Ebf1*-deficient BM niche reduces MF progression. **a.** Experimental setup. **b.** Frequency of MPL^W515L^ mutant (GFP^+^) cells in bone marrow (BM), spleen (SPL) and peripheral blood (PB) of *Prx1*^Cre^*Ebf1*^fl/fl^ and control recipients transplanted with MPL^W515L^ cells. **c.** Spleen weights and representative image of spleens of the recipient mice shown in (b). Each dot represents an individual recipient, data represents the mean ± SEM, *n*=6-8 controls and *n*=5-8 *Prx1*^Cre^*Ebf1*^fl/fl^ mice in 3 independent experiments. **d.** Fibrosis grade scored based on reticulin staining and representative images of reticulin staining in BM sections of mice transplanted with MPL^W515L^ expressing cells. **e.** Bone marrow cellularity of control and *LepR*^Cre^*Ebf1*^fl/fl^ mice transplanted with MPL^W515L^ expressing cells. Frequency of MPL mutant granulocytes (Mac1Gr1^hi^-GFP^+^) and monocytes (Mac1Gr1^lo^-GFP^+^) in BM **(f)**, SPL **(g)**, and PB **(h)** of control and *LepR*^Cre^*Ebf1*^fl/fl^ mice transplanted with MPL^W515L^ cells. Each dot represents an individual recipient, data represents the mean ± SEM, *n*=11-13 controls and *n*=12-15 *LepR*^Cre^*Ebf1*^fl/fl^ mice in 3 independent experiments. **i.** Fibrosis grade based on reticulin staining and representative images of reticulin staining in BM sections of control and *LepR*^Cre^*Ebf1*^fl/fl^ mice transplanted with MPL^W515L^ expressing cells (for Fig. 2d).

**Extended Data Fig. 3.**
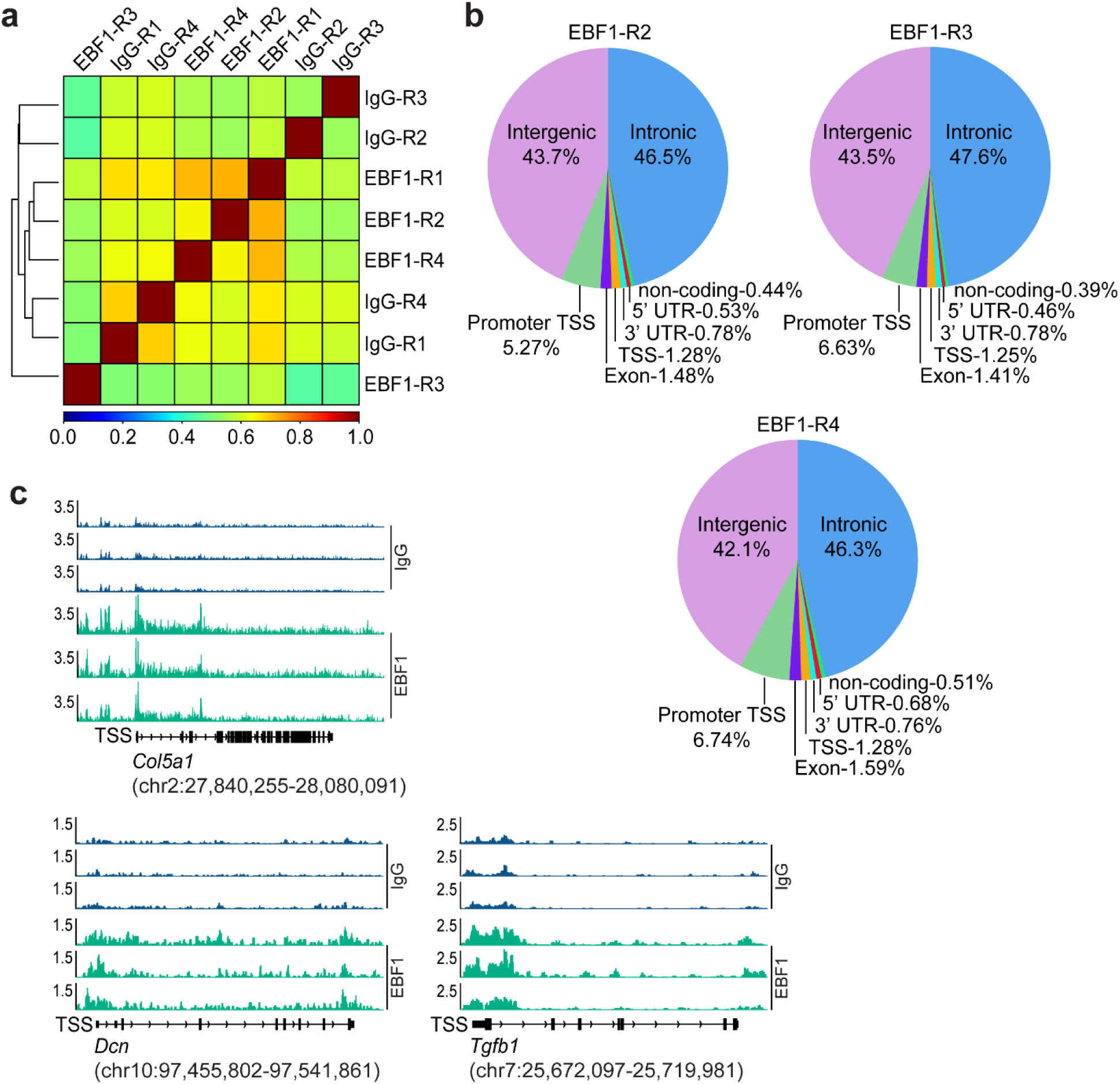
EBF1 chromatin occupancy in fibrotic markers. **a.** Correlation matrix showing hierarchical clustering of independent biological replicates in CUT&RUN analysis assessing EBF1 chromatin occupancy in mouse MSCs. **b.** Distribution of EBF1 peaks in different genomic regions in independent mouse MSCs replicates. **c.** Representative tracks showing EBF1 occupancy within *Col5a1*, *Dcn* and *Tgfb1* loci in mouse MSCs (three independent biological replicates). The scale on the *y* axis represents CPM (count per million) values; transcriptional start site (TSS).

**Extended Data Fig. 4.**
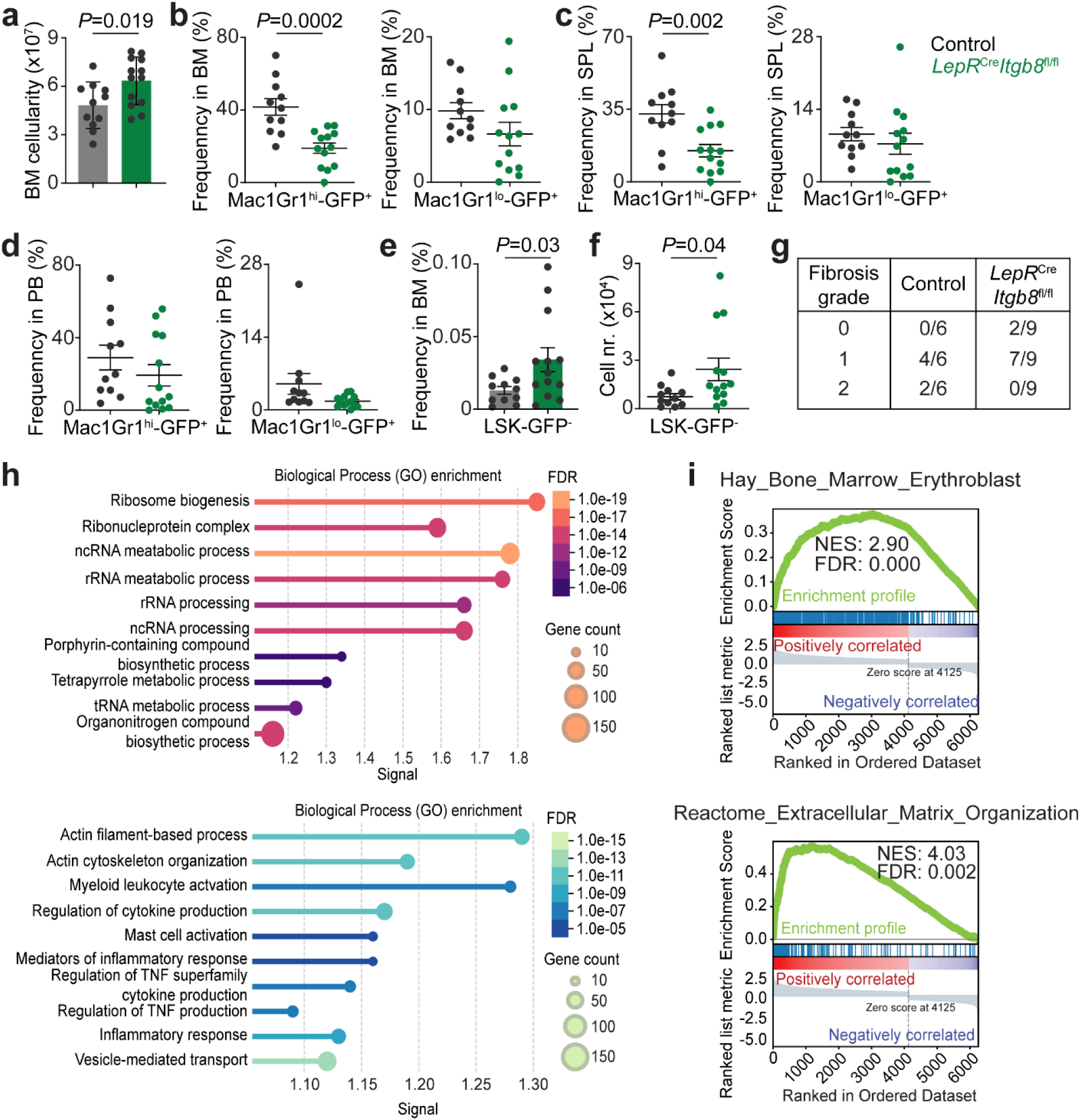
*Itgb8* deficiency in LepR^+^MSCs diminishes MF development. **a.** Bone marrow cellularity of control and *LepR*^Cre^*Itgb8*^fl/fl^ recipients transplanted with MPL^W515L^ expressing cells. Frequency of MPL mutant granulocytes (Mac1Gr1^hi^-GFP^+^) and monocytes (Mac1Gr1^lo^-GFP^+^) in BM **(b)**, SPL **(c)**, and PB **(d)** of control and *LepR*^Cre^*Itgb8*^fl/fl^ mice transplanted with MPL^W515L^ cells. **e.** Frequency and **f.** absolute numbers of non-mutant hematopoietic progenitors (LSK-GFP^−^ cells, Lineage^−^Sca1^+^c-kit^+^) in the BM of control and *LepR*^Cre^*Itgb8*^fl/fl^ recipients transplanted with MPL^W515L^ expressing cells. Each dot represents an individual recipient, data represents the mean ± SEM, *n*=11 controls and *n*=13 *LepR*^Cre^*Itgb8*^fl/fl^ mice in 3 independent experiments. **g.** Fibrosis grade based on reticulin staining and representative images of reticulin staining in BM sections of control and *LepR*^Cre^*Itgb8*^fl/fl^ transplanted with MPL^W515L^ expressing cells (for Fig. 5d). **h.** Gene Ontology (GO) analysis of the upregulated genes (upper panel) and downregulated genes (lower panel) in *Itgb8*-deficient vs control LepR^+^MSCs purified from *LepR*^Cre^*Itgb8*^fl/fl^ and control mice transplanted with MPL^W515L^ cells. **i.** Gene set enrichment analysis (GSEA) from *Itgb8*-deficient vs control LepR^+^MSCs purified from *LepR*^Cre^*Itgb8*^fl/fl^ and control mice transplanted with MPL^W515L^ cells. Normalized Enrichment Score (NES), False Discovery Rate (FDR).

**Extended Data Fig. 5.**
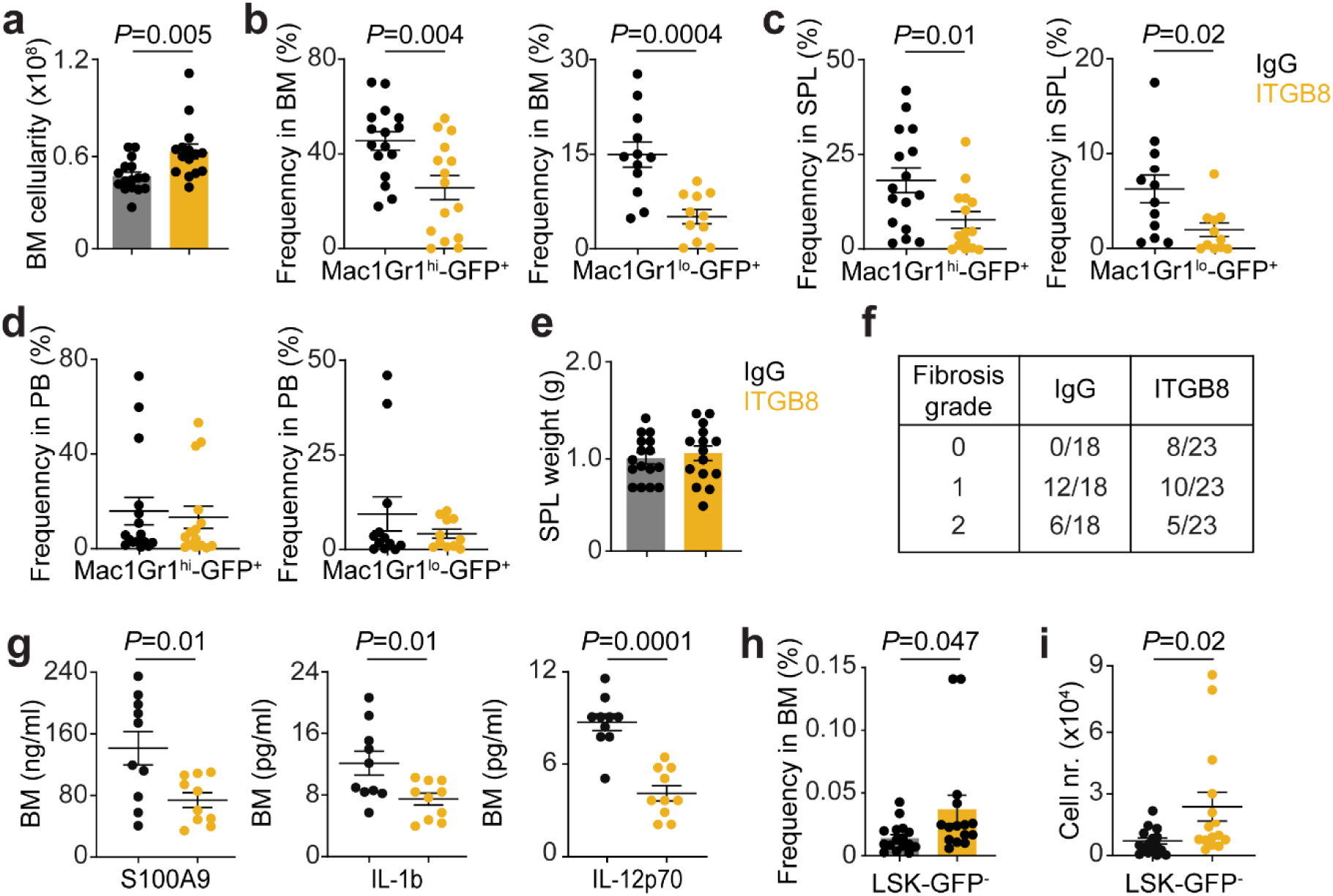
Pharmacological ITGB8 inhibition reduces MF progression. **a.** Bone marrow cellularity of mice transplanted with MPL^W515L^ expressing cells and treated with control IgG or neutralizing antibodies against ITGB8. Frequency of MPL mutant granulocytes (Mac1Gr1^hi^-GFP^+^) and monocytes (Mac1Gr1^lo^-GFP^+^) in BM **(b)**, SPL **(c)**, and PB **(d)** of recipients transplanted with MPL^W515L^ cells and treated with IgGs or anti-ITGB8 antibodies. **e.** Spleen weights of recipients transplanted with MPL^W515L^ cells and treated with IgGs or anti-ITGB8 antibodies. f. Fibrosis grade based on reticulin staining and representative images of reticulin staining in BM sections of mice transplanted with MPL^W515L^ expressing cells and treated with control IgG or anti-ITGB8 antibodies (for Fig. 6d). **g.** Cytokine levels quantified by multiplex cytokine array (Luminex) in the BM supernatants isolated from mice transplanted with MPL^W515L^ cells and treated with control IgG or ITGB8 neutralizing antibodies. Each dot represents an individual recipient, data represents the mean ± SEM, *n*=10 IgG treated and *n*=8-10 anti-ITGB8 treated mice. **h.** Frequency and **i.** absolute numbers of non-mutant hematopoietic progenitors (LSK-GFP^−^ cells, Lineage^−^Sca1^+^c-kit^+^) in the BM of transplanted with MPL^W515L^ cells and treated with IgGs or anti-ITGB8 antibodies. Each dot represents an individual recipient, data represents the mean ± SEM, *n*=16 IgG treated and *n*=15 anti-ITGB8 treated mice.

**Extended Data Fig. 6.**
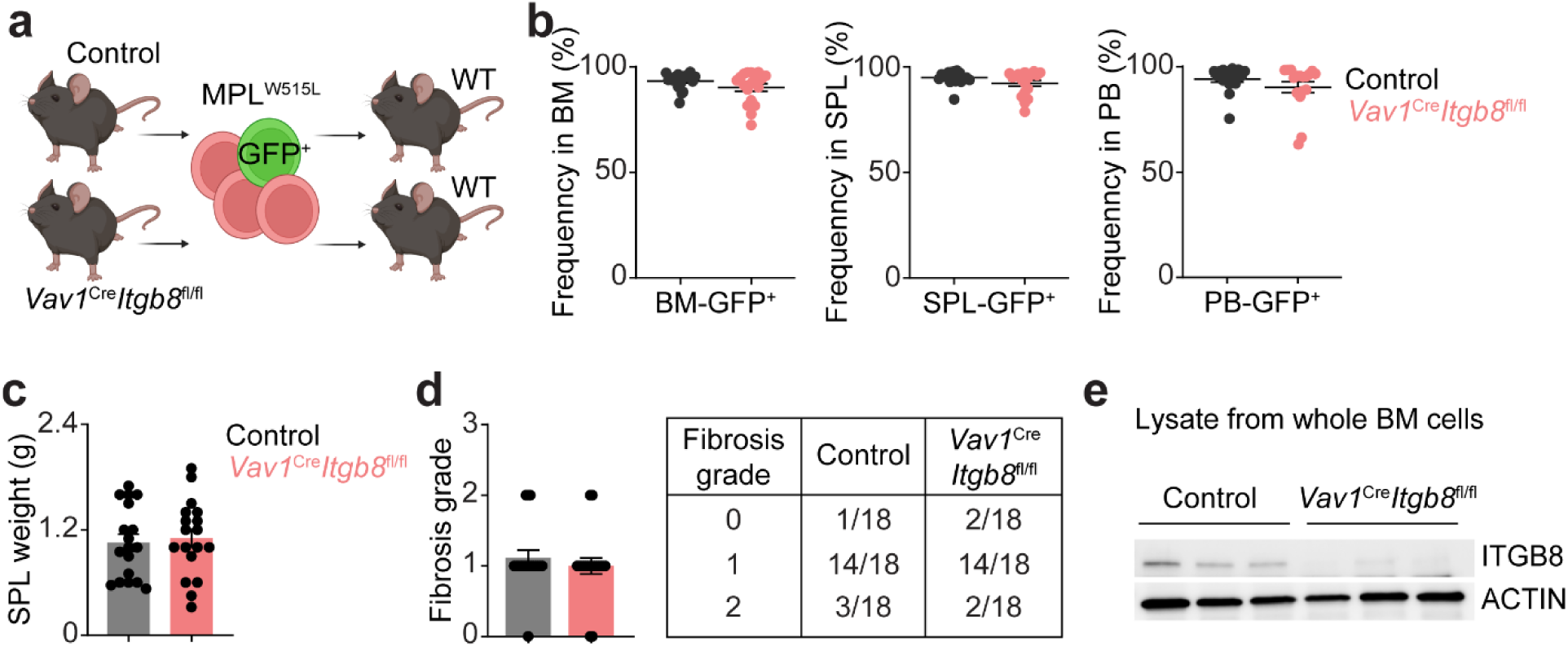
*Itgb8* deletion in hematopoietic cells does not affect MF development. **a.** Experimental setup. **b.** Frequency of MPL^W515L^ mutant (GFP^+^) cells in the BM, SPL, and PB of wild-type recipients transplanted with c-kit^+^ cells isolated from control and *Vav1*^Cre^*Itgb8*^fl/fl^ mice and transduced with MPL^W515L^ construct. **c.** Spleen weights of the recipients shown in (b). Each dot represents an individual recipient, data represents the mean ± SEM, *n*=17-18 recipients of c-kit^+^ cells isolated from 8 control mice and transduced with MPL^W515L^ and *n*=17-18 recipients of c-kit^+^ cells isolated from 9 *Vav1*^Cre^*Itgb8*^fl/fl^ mice and transduced with MPL^W515L^ in 3 independent experiments. **d.** Fibrosis grade scored based on reticulin staining of BM sections from wild-type recipients transplanted with c-kit^+^ cells isolated from control and *Vav1*^Cre^*Itgb8*^fl/fl^ mice and transduced with MPL^W515L^ construct. **e.** Western blot analysis showing ITGB8 protein level in BM cells from control and *Vav1*^Cre^*Itgb8*^fl/fl^ mice.

